# Nonlinear Elastic Bottlebrush Polymer Hydrogels Modulate Actomyosin Mediated Protrusion Formation in Mesenchymal Stromal Cells

**DOI:** 10.1101/2024.03.08.584195

**Authors:** Monica L. Ohnsorg, Kayla M. Mash, Alex Khang, Varsha V. Rao, Bruce E. Kirkpatrick, Kaustav Bera, Kristi S. Anseth

## Abstract

The nonlinear elasticity of many tissue-specific extracellular matrices is difficult to recapitulate without the use of fibrous architectures, which couple strain-stiffening with stress relaxation. Herein, bottlebrush polymers are synthesized and crosslinked to form poly(ethylene glycol)-based hydrogels and used to study how strain-stiffening behavior affects human mesenchymal stromal cells (hMSCs). By tailoring the bottlebrush polymer length, the critical stress associated with the onset of network stiffening is systematically varied, and a unique protrusion-rich hMSC morphology emerges only at critical stresses within a biologically accessible stress regime. Local cell-matrix interactions are quantified using 3D traction force microscopy and small molecule inhibitors are used to identify cellular machinery that plays a critical role in hMSC mechanosensing of the engineered, strain-stiffening microenvironment. Collectively, this study demonstrates how covalently crosslinked bottlebrush polymer hydrogels can recapitulate strain-stiffening biomechanical cues at biologically relevant stresses and be used to probe how nonlinear elastic matrix properties regulate cellular processes.

## MAIN

Biological tissues are characterized by a balance of viscoelastic and nonlinear elastic properties that depend on the tissue specific extracellular matrix (ECM) and its interactions with resident cells^[1,2]^. To date, viscoelastic hydrogels have been widely used to study cell mechanosensing, migration, differentiation, and other functional properties which have been extensively reviewed^[3,4]^. While these synthetic ECM mimics can be tailored with biologically relevant material properties including stress relaxation over a wide range of timescales, it has been difficult to engineer synthetic hydrogels with the complementary strain-stiffening response. Nonlinear elastic behavior, specifically strain-stiffening, is typically observed in proteinaceous materials such as collagen and fibrin^[1]^. These ECM macromolecules in gel form exhibit strain-stiffening as isotropic semi-flexible filamentous networks^[5]^. However, such fibrous materials exhibit nonlinear viscoelasticity – an interplay of nonlinear elasticity over short timescales and viscoelastic dissipation of stresses over longer timescales – which confounds the isolated effect of nonlinear elastic biomechanical cues^[2]^.

Strain-stiffening mechanical properties can be accessed by any network composed of semiflexible filaments, but it has yet to be seen if the synthetic origin of strain-stiffening mechanics affects cell behavior^[6]^. Bottlebrush polymers, consisting of a linear polymer backbone from which polymer side chains emanate radially, capture the rheological properties of molecular filaments^[7]^. Recently, bottlebrush polymers have been incorporated into network structures to form solvent free elastomers^[8,9]^, plastomers^[10,11]^, sensors^[12,13]^, and adhesives^[14–16]^. Sheiko, Dobrynin, and coworkers used linear-bottlebrush-linear (LBL) triblock copolymers that undergo microphase separation to form composite networks and observed bottlebrush polymers to behave as semiflexible filaments between microphase-separated crosslinking points, yielding nonlinear elastic material properties under elastic deformation^[10,17– 19]^. These results demonstrated that LBL materials can be used to form plastomers and hydrogels that mimic nonlinear elastic bulk tissue mechanics^[11,17,20]^. However, bottlebrush hydrogels have yet to be utilized as a three-dimensional (3D) cell culture platform to mimic and isolate strain-stiffening biomechanical properties. Therefore, in this work we adapt the LBL strain-adaptive network architecture for temperature independent photo-crosslinking using norbornene-functionalized 8-arm star-PEG macromers connected by dithiol bottlebrush polymers. In doing so, we access strain-stiffening hydrogels with well-defined network connectivity and facile incorporation of cell adhesion peptides.

To date, synthetic strain-stiffening hydrogels used for 3D culture have been largely fibrous in architecture^[21–24]^. For example, Rowan and coworkers synthesized a thermoresponsive polyisocyanopeptide (PIC) hydrogel composed of hydrogen-bond stabilized β-helical polymer backbones which assemble into fibrillar bundles^[21,25]^. This synthetic filamentous hydrogel mimics the non-linear elastic properties of proteinaceous hydrogels and can be molecularly tuned to achieve a variety of moduli and stiffening profiles^[1]^. Consequentially, these materials access stress-stiffening at critical stresses (σ_*c*_) within the biologically relevant stress regime (BRSR), which has been loosely defined as where filamentous ECM components – such as collagen, fibrin, and actin – stiffen (σ_*c*_< 25 Pa, γ_*c*_< 10%)^[5]^. Using these materials, the differentiation potential and spreading of human adipose-derived stem cells could be directed as a function of σ_*c*_^[8,26]^. However, the cellular machinery and mechanisms responsible for sensing strain-stiffening microenvironments remain unknown. It is also still unclear what downstream pathways regulate cell phenotype changes as a function of σ_*c*_.

Herein, we engineered bottlebrush polymer hydrogel networks for 3D cell culture and used bone marrow-derived human mesenchymal stem cells (hMSCs) to systematically assess the effects of σ_*c*_ on hMSC morphology and mechanotransduction around the BRSR. hMSCs were selected as the model cell type since they are mechanically responsive and reside in the collagenous strain-stiffening osteoid *in vivo*. We characterized cell morphology as a function of σ_*c*_ and inspected how the matrix dictated cell behavior in response to stiffening irrespective of bulk modulus. When cultured within bottlebrush hydrogels with σ_*c*_ within the BRSR, hMSCs formed focal adhesion-mediated protrusions as a function of σ_*c*_. The forces exerted by the cells within these bottlebrush networks were visualized and quantified using 3D traction force microscopy. Actomyosin dynamics governing the hMSC response to strain-stiffening were investigated using small molecule inhibitors to block myosin II adenosine triphosphatase activity and stabilize the inactive state of Arp2/3 complex to inhibit cell contractility and actin branching, respectively. This suppressed protrusion formation and identified the cellular machinery involved in strain-stiffening mechanotransduction. By decoupling the variable of nonlinear elasticity from a fibrous architecture using a covalently crosslinked bottlebrush polymer hydrogel, one can elucidate critical features of this biomechanical feedback mechanism on a cellular level and better understand its impact in native ECM and living tissues.

### Bottlebrush Polymer Hydrogels with Tunable Levels of Strain-Stiffening

A difunctional trithiocarbonate chain transfer agent was used to polymerize poly(ethylene glycol) monomethyl ether acrylate (PEGA9, average *M*n = 480 Da, *N*sc = 9) via reversible-addition fragmentation chain transfer techniques to synthesize four C12H25-PPEGA9-*N*bb-C12H25 bottlebrush macromers with increasing backbone degree of polymerization, *N*bb (**Figure 1a**, **Table 1**, Supplementary Information Section 1). The trithiocarbonate end-groups were then cleaved to free thiols via aminolysis, yielding four bottlebrush dithiol macromomers (SH-PPEGA9-*N*bb-SH) with *N*bb of 20, 50, 80, and 110 (**Figure 1b**, **Figure S7**). These macromers were crosslinked into a network with an 8-arm, 20 kDa PEG-norbornene (PEG-NB) using 5 mW/cm^2^ of 365 nm light for 5 minutes in the presence of photoinitiator (LAP, 6.8 mM) and cell-adhesion peptide (CRGDS, 1 mM) (**Table S2**). As a control, the bottlebrush crosslinked networks were compared to modulus-matched hydrogels formed using linear 2 kDa PEG-dithiols (SH-PEG-2k-SH) as crosslinkers at two different weight percents (PEG-2k, **3.5wt%** and PEG-2k, **2.5wt%**). The SH-PEG-2k-SH was selected for this study to reduce network entanglements, as the entanglement molecular weight (*M*e) of PEG is 4,400 Da^[27]^. In doing so, we achieved similar network connectivity across formulations since the bottlebrush polymer *M*e is >100 kDa^[28]^ (**Fig 1c-d**).

**Figure 1.**
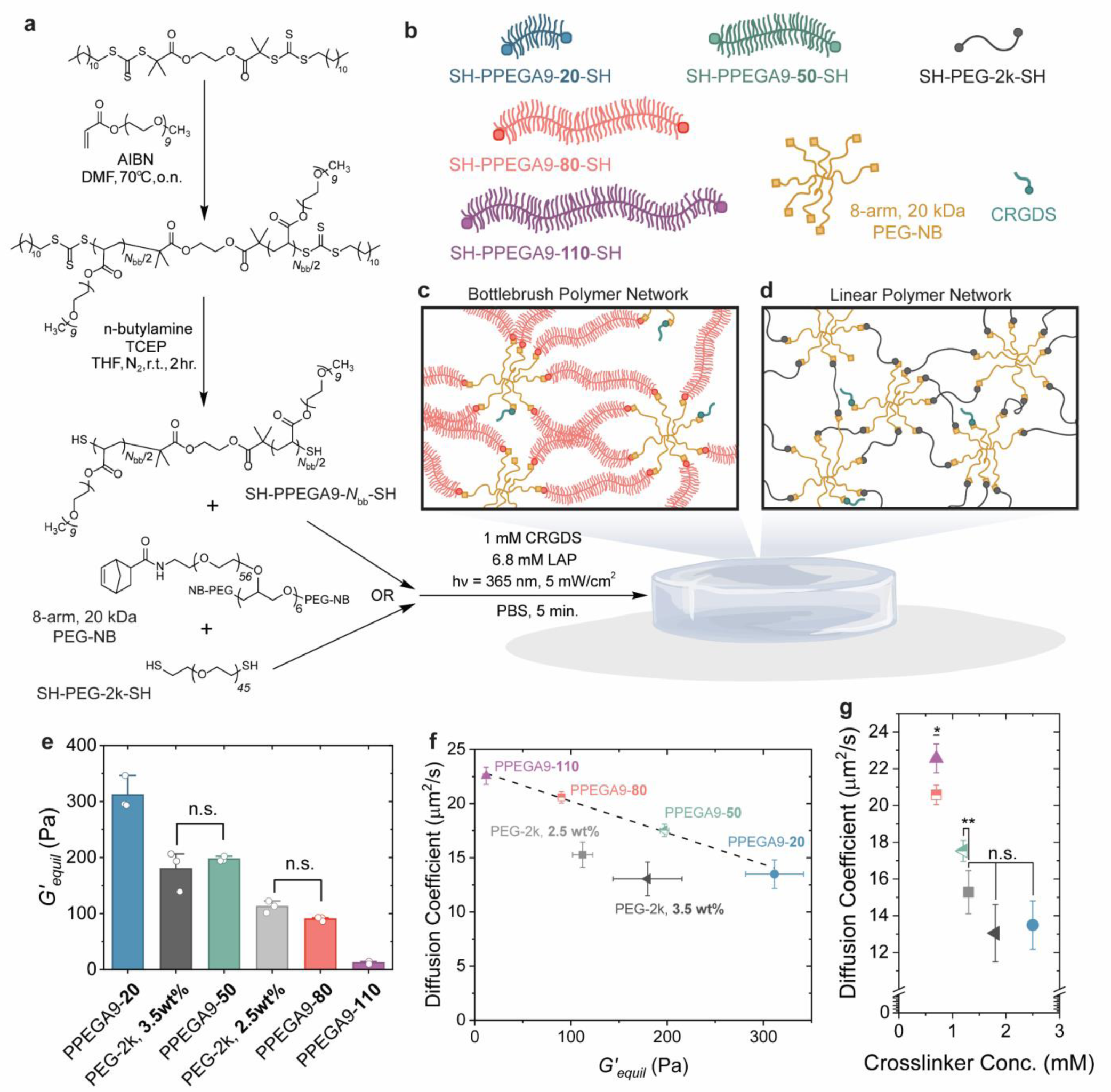
Synthesis and characterization of photocrosslinked bottlebrush polymer hydrogels. **a,** Synthetic scheme of the bottlebrush dithiol synthesis and subsequent photocrosslinking with an 8-arm 20 kDa star PEG-norbornene. The 2 kDa linear PEG dithiol hydrogel formulation is also shown. **b,** Illustrated representation of the bottlebrush polymer dithiols, PEG-2k dithiol, 8-arm PEG-NB crosslinker, and CRGDS. **c,** Illustration of bottlebrush polymer network architecture. **d,** Illustration of linear polymer network architecture. **e,** Equilibrium swollen moduli measured for each hydrogel formulation ordered from highest to lowest modulus (n=3). **f,** Diffusion coefficient of FITC-BSA within each hydrogel formulation measured using fluorescence recovery after photobleaching (FRAP) and plotted against the equilibrium swollen storage modulus (*G’_equil_*). The bottlebrush polymer hydrogels had higher diffusion coefficients than the linear PEG hydrogels (dotted line, r^2^ = 0.99). **g,** Plot of diffusion coefficient vs. concentration of 8-arm PEG-NB crosslinker in each formulation. The PPEGA9-**20** (blue circle), PEG-2k, **3.5wt%** (grey triangle), and PEG-2k, **2.5wt%** (grey square) each had similar diffusion coefficients despite the PPEGA9-**20** incorporating more crosslinks. PPEGA9-**50** (green, half-filled triangle) had a greater diffusion coefficient than the PEG-2k, **2.5wt%** sample with a similar crosslinker concentration. In addition, the longest bottlebrush samples had the greatest diffusion coefficients and the PPEGA9-**110** (purple triangle) had a greater measured diffusion coefficient than PPEGA9-**80** (salmon, half-filled square) despite having the same concentration of crosslinker.

**Table 1.** Summary of Hydrogel Formulations and Mechanical Properties.

| Dithiol Macromer | $M_{n, SEC}^a$<br>[kDa] | SH:NB | Initial<br>wt% | Dithiol<br>conc.<br>[mM] | 8-arm<br>20 kDa<br>PEG-NB<br>conc.<br>[mM] | $G'_{equil}^b$<br>[Pa] | $\sigma_{c, equil}^c$<br>[Pa] | $m^d$ |
| --- | --- | --- | --- | --- | --- | --- | --- | --- |
| SH-PPEGA9-**20**-SH | 9.8 | 1.0:1.0 | 10.0 | 9.6 | 2.5 | 311 ± 30 | 76 ± 7.4 | 1.5 ± 0.1 |
| SH-PEG-2k-SH | 2.0 | 1.0:1.0 | 3.5 | 6.8 | 1.8 | 180 ± 36 | 49.3 ± 28.6 | 1.5 ± 0.2 |
| SH-PPEGA9-**50**-SH | 24.5 | 1.0:1.0 | 10.0 | 4.9 | 1.2 | 197 ± 5 | 36.5 ± 9.7 | 1.5 ± 0.1 |
| SH-PEG-2k-SH | 2.0 | 1.0:1.0 | 2.5 | 4.9 | 1.3 | 112 ± 11 | 27.3 ± 2.6 | 1.5 ± 0.01 |
| SH-PPEGA9-**80**-SH | 40.2 | 1.0:1.0 | 10.0 | 3.2 | 0.7 | 90 ± 4 | 21.6 ± 0.8 | 2.0 ± 0.02 |
| SH-PPEGA9-**110**-SH | 55.3 | 1.0:1.2 | 10.5 | 2.5 | 0.7 | 12 ± 3 | 2.0 ± 0.4 | 2.8 ± 0.1 |

<sup>a</sup>) SEC-MALS in DMF with 0.05M LiBr. <sup>b</sup>) Shear modulus measured at 1 % strain, and 1.0 rad/s with an axial force of 0.1 N. <sup>c</sup>) Scaled *equilibrium swollen* critical stress based on the *in situ* critical stress measurements. The critical stress was determined using a knee-elbow analysis of each $K'$ vs. critical stress plot<sup>[29]</sup>. <sup>d</sup>) Stiffening index calculated from the slope of the line past the critical stress.

In all systems, network evolution was monitored by oscillatory shear rheometry and the G”/G’ crossover was used as an estimate of the gel point (**Figure S9**, Table **S2**). The storage modulus of each hydrogel was measured after equilibrium swelling, *G’equil,* to best represent the mechanical microenvironment during cell culture (**Table 1**). The bottlebrush polymer hydrogels decreased in *G’equil* with increasing *N*bb, consistent with increasing mesh size. The PPEGA9-**50** was modulus-matched to PEG-2k, **3.5wt%** (*G’equil* ≈ 200 Pa), and storage modulus of PPEGA9-**80** was not significantly different from PEG-2k, **2.5wt%** (*G’equil* ≈ 100 Pa) (**Figure 1e**, **Table 1**). Using fluorescence recovery after photobleaching (FRAP) to measure the diffusion of FITC-BSA in equilibrated networks, we observed a higher diffusivity in the bottlebrush hydrogels compared to the moduli-matched linear networks (**Fig 1f**), reinforcing that bottlebrush dithiols act as semiflexible filaments separating network junctions. This phenomenon can be visualized in **Figure 1g** where the PPEGA9-**80** and PPEGA9-**110** formulations have the same initial concentration of 8-arm 20 kDa PEG-NB, but the measured diffusion coefficient for the PPEGA9-**110** hydrogel is larger due to the 30 additional repeat units in the bottlebrush *N*bb extending the distance between crosslinks (Table **S2**). Of note, these bottlebrush polymer hydrogels afford access to ultrasoft moduli similar to proteinaceous hydrogels (e.g. fibrin and collagen) while also recapitulating their strain-stiffening mechanical properties (**Figure S10**).

The strain-stiffening mechanics of the hydrogels were measured directly after network formation (*equilibrium*/*in situ* scaling, Table S3) and plotted as oscillation stress versus strain (**Figure 2a**). The strain-stiffening properties of the bottlebrush scaffolds can be attributed to the high *M*_e_ of the SH-PPEGA9-***N*_bb_**-SH, which only need to be stretched a small amount to reach complete network extension and the onset of stiffening compared to linear PEG dithiols as illustrated in **Figure 2b**^[30]^. When the differential modulus, *K’equil*, is plotted as a function of σ_*equil*_ to visualize the onset of strain-stiffening (σ_*c*,*equil*_), we observe that σ_*c*,*equil*_ of the bottlebrush hydrogels is always lower than the moduli-matched PEG-2k, **2.5** and **3.5 wt%** controls. (Table 1, **Fig 2c-d**). The σ_*c*,*equil*_ scaled linearly with *G’equil* (r^2^ = 0.98) (**Figure 2d**, S11) similar to previously characterized fibrous PIC^[26]^ and EKGel^[31]^ systems. While fibrous hydrogels follow an affine model of network deformation by rigid filaments^[31]^, the critical strain of the covalently crosslinked structures studied here remained consistent across formulations at 40% strain (**Figure S12**). The measured σ_*c*,*equil*_ for both PPEGA9-**80** (22 Pa) and PPEGA9-**110** (2 Pa) indicates stiffening within the BRSR (< 25 Pa). The σ_*c*,*equil*_ of the modulus-matched PEG-2k, **2.5wt%** was just outside the BRSR (27 Pa). Therefore, the bottlebrush polymer networks facilitate the ability to access σ_*c*,*equil*_ within the BRSR using a non-fibrous, elastic (**Figure S13**) hydrogel. The stiffening index, *m*, of each formulation was measured as the slope of the line after σ_*c*,*equil*_. In “high σ_*c*,*equil*_” samples (PPEGA9-**20**, PEG-2k, **3.5wt%**, and PPEGA9-**50**), *m* was 1.5; however, “low σ_*c*,*equil*_” samples had *m* between 1.9 to 2.8 (**Table 1**, **Fig 2d**), indicating they are more responsive to applied stresses. Previously observed *m* values for PIC hydrogel systems have all been ≤ 1.5^[25,26]^. Therefore, isotropic, nanoporous networks have the potential to stiffen at greater rates than synthetic fibrous networks, thus allowing bottlebrush networks to recapitulate the mechanics of proteinaceous hydrogel matrices more faithfully (**Figure S10**).

**Figure 2.**
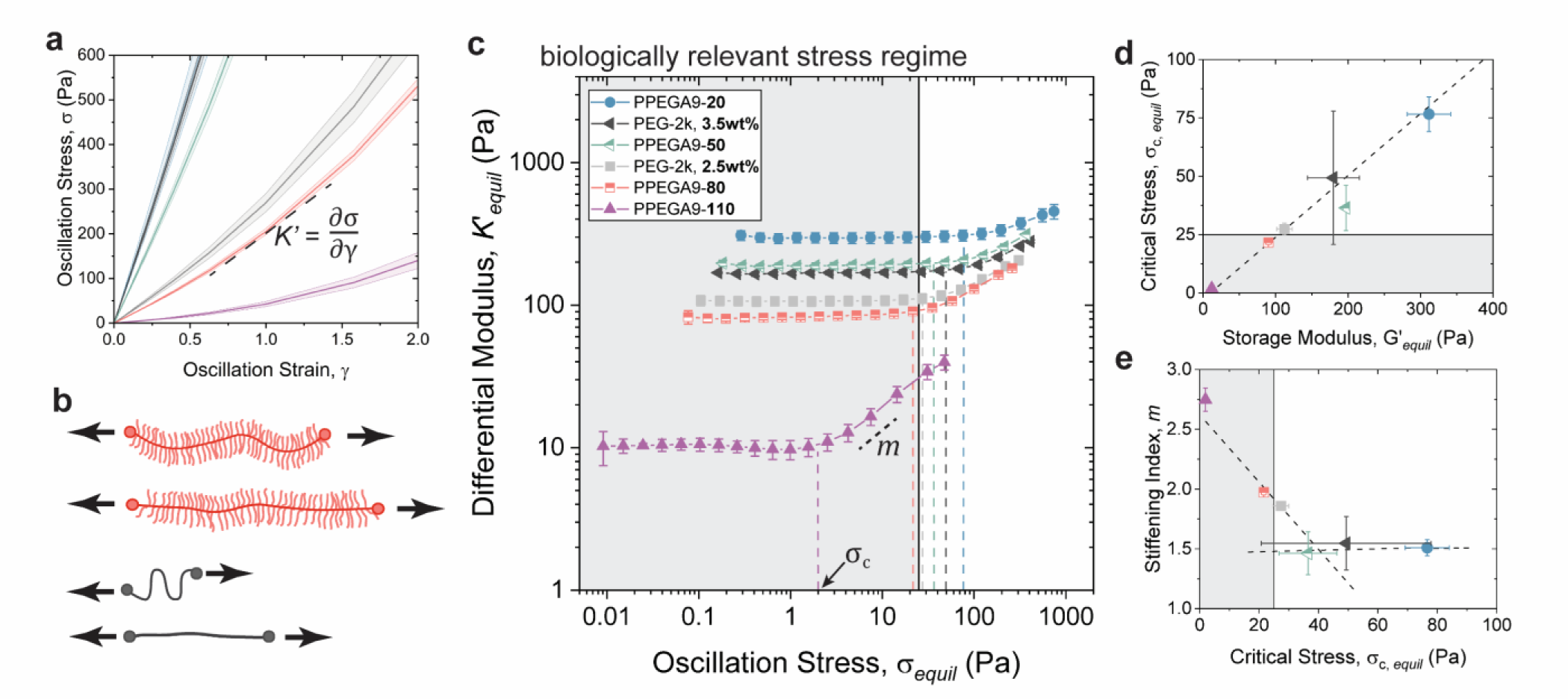
Strain-stiffening in covalently crosslinked hydrogels. **a,** Measured oscillation stress plotted as a function of oscillation strain for each hydrogel formulation. (PPEGA9-**20**, blue; PEG-2k, **3.5 wt%**, dark grey; PPEGA9-**50**, green; PEG-2k, **2.5wt%**, light grey; PPEGA9-**80**, salmon; PPEGA9-**110**, purple) **b,** Illustration of the deformation levels needed to fully extend a bottlebrush dithiol crosslinker as opposed to a linear dithiol crosslinker. **c**, Equilibrium differential modulus, *K*’*_equil_*, plotted versus equilibrium oscillation stress applied to all six hydrogel formulations. The critical stress, σ_*c*,*equil*_, is denoted as the onset of stiffening (dotted line) and the stiffening index, *m*, is the rate of stiffening after the σ_*c*,*equil*_. (n=3 independent measurements) **d,** σ_*c*,*equil*_ plotted as a function of G’*_equil_*showing that the two parameters are linearly correlated (r^2^ = 0.98, dotted line) and highlighting how the two longest bottlebrush formulations achieve σ_*c*,*equil*_ within the biologically relevant stress regime (<25 Pa). **e,** The average stiffening index, *m*, plotted as a function of σ_*c*,*equil*_ with two independent fit lines showing no linear correlation at high σ_*c*,*equil*_ (r^2^ = 0.23, dotted line) and a strong linear correlation at low σ_*c*,*equil*_ (r^2^ = 0.93, dotted line).

### Biologically accessible nonlinear elasticity influences hMSC mechanosensing and morphology

To investigate the cellular response to the transition of strain-stiffening behavior from outside to within the BRSR, we encapsulated hMSCs within the six hydrogel formulations. In all hydrogel formulations, the cells remained viable over six days (**Figure S14-15**). The hMSCs exhibited a spherical morphology in the PPEGA9-**20**, PEG-2k, **3.5 wt%**, and PPEGA9-**50** hydrogels, which all had σ_*c*,*equil*_ >35 Pa (**Figure 3a,i-iii**). This spherical morphology is typical for cells encapsulated in non-degradable, elastic hydrogels, where spreading is constrained by the nanoporous network structure^[32]^. However, at σ_*c*,*equil*_< 30 Pa (**Figure 3aiv-vi**), we observe a transition from primarily spherical cellular morphologies just outside the BRSR – PEG-2k, **2.5wt%** formulation (σ_*c*,*equil*_ = 27 Pa) – to the initial appearance of spindle-like protrusions in PPEGA9-**80** formulation just within the BRSR (σ_*c*,*equil*_ = 22 Pa). A similar protrusive phenotype for human adipose-derived stem cells and fibroblasts has been observed within synthetic and natural fibrous hydrogels, which exhibit strain-stiffening within the BRSR^[26,33,34]^. However, these previous findings are confounded by the propensity of fibrous materials to also stress-relax, making it difficult to definitively determine the impact of strain-stiffening mechanics on cell behavior. Therefore, cell culture in these complementary non-degradable, elastic, and strain-stiffening bottlebrush hydrogels aids in identifying cell phenotypes that uniquely depend on nonlinear elastic material properties.

**Figure 3.**
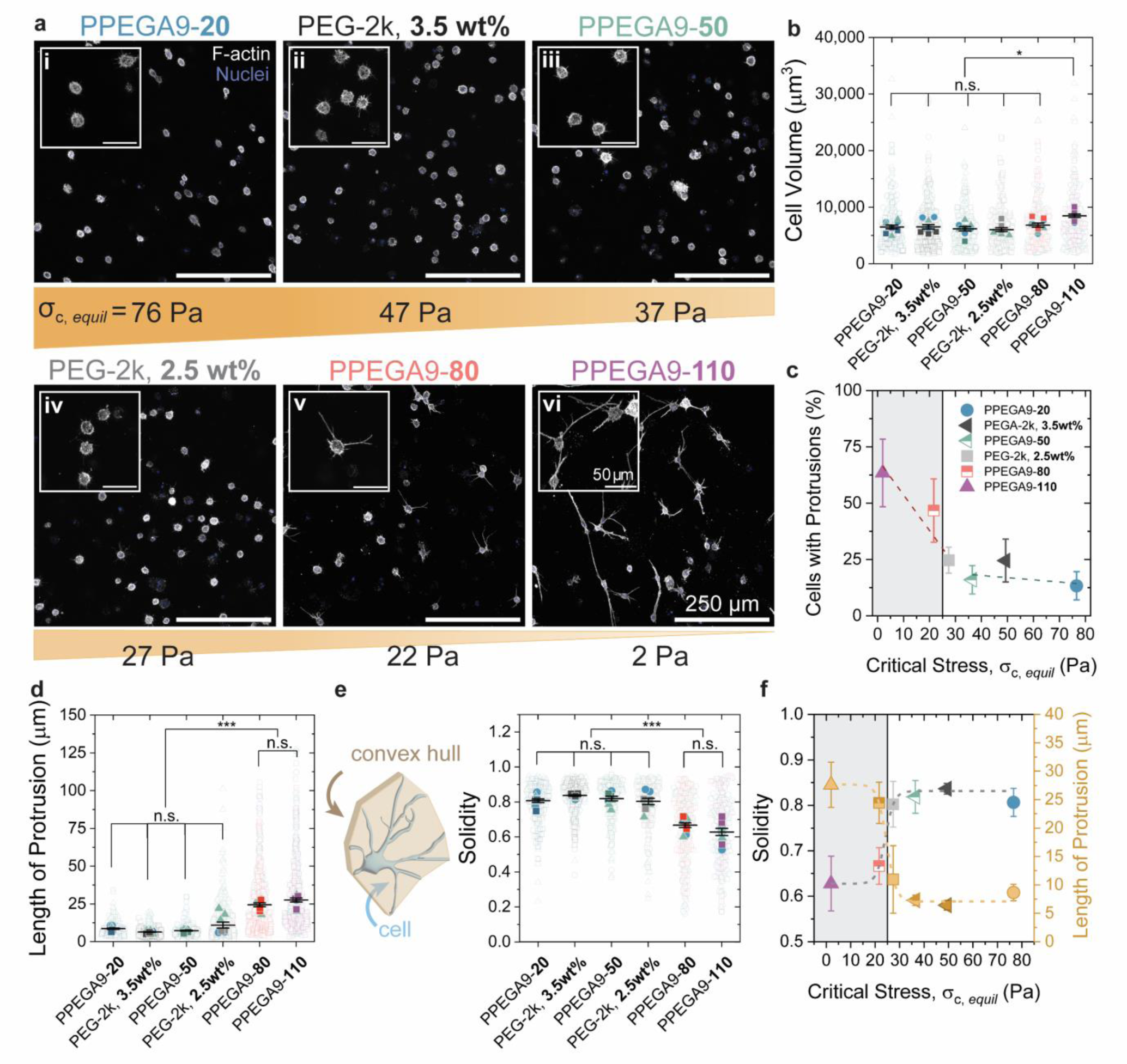
hMSC morphology and mechanotransduction as a function of nonlinear elasticity. **a,** Representative confocal images of hMSCs encapsulated in each hydrogel formulation fixed at day 3 and stained for DAPI (blue) and f-actin (grey). (i) PPEGA9-**20**, (ii) PEG-2k, **3.5wt%**, (iii) PPEGA9-**50**, (iv) PEG-2k, **2.5wt%**, (v) PPEGA9-**80**, and (vi) PPEGA9-**110** are ordered from highest to lowest σ_*c*,*equil*_. **b**, Super plot showing the average cell volume across hydrogel conditions. **c,** The average % of cells with protrusions plotted as a function of critical stress, σ_*c*,*equil*_, with two independent fit lines showing no linear correlation at high σ_*c*,*equil*_ (r^2^ = 0.20) and a strong linear correlation (slope = -1.6 ± 0.5, r^2^ = 0.89) at low σ_*c*,*equil*_. **d,** Super plot of the length of each protrusion. **e,** Illustration of the convex hull surrounding a cell needed to calculate the solidity. Super plot denoting the solidity measurements of hMSCs encapsulated within each hydrogel formulation. **f,** Solidity and length of protrusions plotted as a function of σ_*c*,*equil*_. Dotted lines are Boltzmann fits of the data and are present to guide the eye indicating the inversion of properties within and outside the biologically relevant stress regime. (n= 3 independent biological replicates)

The transition towards the protrusive morphology as a function of σ_*c*,*equil*_ was quantified using cell volume and protrusion metrics. Protrusions around generally spherical cell bodies did not significantly increase average cell volume, but cell volume did increase in PPEGA9-**110** (σ_*c*,*equil*_ = 2 Pa) hydrogels (**Figure 3b**), in which >50% of the encapsulated hMSCs had protrusions averaging >25 µm in length (**Figure 3c-d**). Of note, the percentage of cells with protrusions did not correlate with the σ_*c*,*equil*_ outside the BRSR; however, within the BRSR, the percentage of cells with protrusions increased as a function of decreasing σ_*c*,*equil*_ (**Figure 3c**). To better characterize hMSC morphology as a function of σ_*c*,*equil*_, the length of each protrusion was measured (**Figure 3d**) and the complementary 3D metric of solidity was calculated. Solidity defines the space-filling nature of each cell by using the ratio of cell volume divided by the volume of its convex hull, the smallest polyhedron which encloses the entire cell (**Figure 3e**). By these metrics, hMSCs cultured in formulations where σ_*c*,*equil*_ was within the BRSR were morphometrically distinct – protrusion length = 25-30 um and solidity < 0.7 – from samples with σ_*c*,*equil*_> 25 Pa (**Figure 3f**).

Given the distinct variation in morphology as a function of σ_*c*,*equil*_, we next investigated the role of cell-matrix interactions in mechanosensing of nonlinear elasticity by quantifying the volume and number of focal adhesions (FAs, paxillin, **Figure 4a**) formed between cells and surrounding matrix. While the average number of FAs did not change across the conditions, the volume of FAs per cell increased in the PPEGA9-**80** and -**110** formulations, where σ_*c*,*equil*_< 25 Pa (**Figure 4b**). Interestingly, the FAs were largest along the cellular protrusions (**Figure 4av-vi**), suggesting that hMSCs may use FAs to probe the surrounding matrix, but only mature upon local stiffening of the surrounding microenvironment. Of further note, cell FAs were larger in all the bottlebrush polymer hydrogels formulations compared to the linear PEG-2k controls (**Figure 4b**).

**Figure 4.**
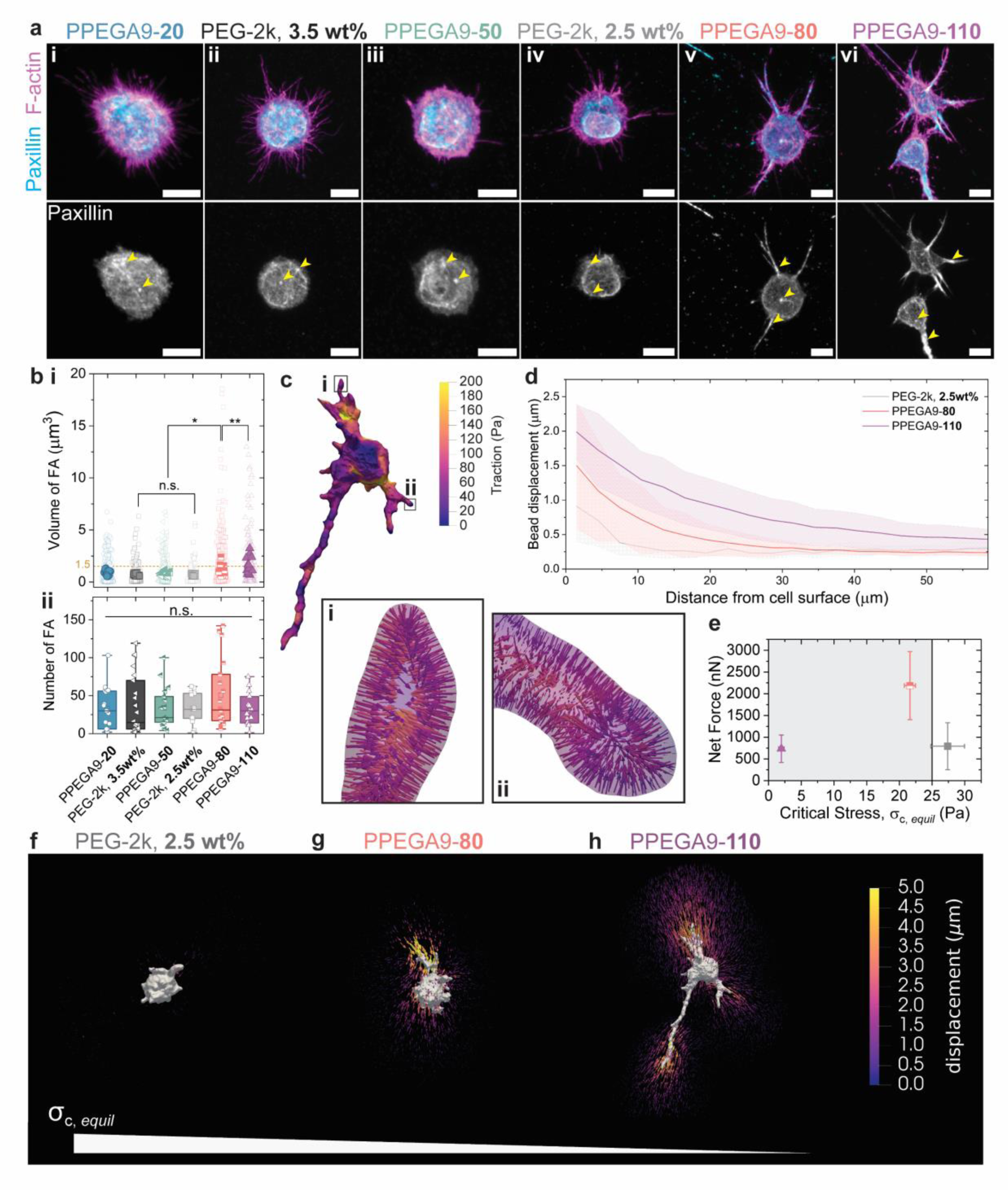
Cell-matrix interactions and cellular displacement of the surrounding elastic hydrogel. **a,** Representative maximum intensity projections of confocal images of hMSCs encapsulated in each hydrogel formulation at Day 3 and immunostained for DAPI (blue), f-actin (magenta), and paxillin (cyan); (i) PPEGA9-**20**, (ii) PEG-2k, **3.5wt%**, (iii) PPEGA9-**50**, (iv) PEG-2k, **2.5wt%**, (v) PPEGA9-**80**, and (vi) PPEGA9-**110** are ordered from highest to lowest σ_*c*,*equil*_. Yellow arrowheads guide the eye to focal adhesions (scale bar = 10 µm) **b**, Quantification of the i) average volume and ii) number of focal adhesions per cell (µm^3^) (n = 10 cells) **c,** Representative image of the traction field across the cell body illustrating the direction and magnitude of each traction vector at the ends of the protrusion (i, ii) **d,** Quantification of bead displacement as a function of distance from the cell surface. **e,** Net force exerted by cells encapsulated within each hydrogel formulation (PEG-2k, **2.5wt%**, grey square; PPEGA9-**80**, salmon half-filled square; PPEGA9-**110**, purple triangle) plotted as a function of σ_*c*,*equil*_. Representative images to visualize the bead displacement vectors for **f,** PEG-2k, **2.5wt%**, **g,** PPEGA9-**80**, and **f,** PPEGA9-**110**. (n = at least 5 cells for each measurement)

To quantify the forces transmitted by hMSC-hydrogel interactions, we used 3D traction force microscopy (3DTFM). Fluorescent beads were embedded within hMSC-laden hydrogel formulations and, following 3 days of culture, the bead positions were tracked before and after treatment with cytochalasin D to disrupt the actin cytoskeleton (**Figure S17**). The traction field along the boundary of each cell was computed from the measured displacement field. A representative traction field is shown for an hMSC encapsulated in the PPEGA9-**110** sample in **Figure 4c**, illustrating how an individual cell can experience varying traction forces along its protrusions versus cell body. The largest bead/hydrogel displacement was observed in the lowest σ_*c*,*equil*_ sample, PPEGA9-**110** (**Figure 4d,h**), where forces propagated further away from the hMSC surface into the matrix. These measurements are consistent with findings related to how cells use nonlinear elastic mechanics to transduce and sense forces over long distances^[35]^. Comparing between the modulus-matched samples, PPEGA9-**80** and PEG-2k, **2.5wt%**, hMSCs displaced the beads much further in the bottlebrush samples with σ_*c*,*equil*_ within the BRSR (**Figure 4d,f-g**). The calculated forces applied by cells encapsulated within the PPEGA9-**80** hydrogel were nearly double those in the modulus-matched PEG-2k, **2.5wt%** formulation (**Figure 4e**). We posit that cells cultured within hydrogels with σ_*c*,*equil*_ within the BRSR have more protrusions and larger focal adhesions that lead to the exertion of greater forces throughout the surrounding material microenvironment. Interestingly, calculated traction forces within the PPEGA9-**110** sample were similar to the PEG-2k, **2.5wt%** condition. We hypothesize that, due to the low σ_*c*,*equil*_ and *G’equil*, cells only need apply a small amount of force to experience a strong strain-stiffening response, eliciting a dramatic change in morphology. However, mechanotransduction in these ultrasoft matrices did not appear to require nuclear translocation of the transcriptional co-activator, YAP, unlike other well-characterized modes of mechanosensing^[36,37]^ (**Figure S18**). Nonlinear elasticity in bottlebrush-derived hydrogels promotes formation of protrusions with mature focal adhesions and leads to generation of traction forces that extend tens of microns into the hydrogel (**Figure 4f-g**). Since bead displacements and traction forces are greatest near hMSC protrusions into the hydrogel, we next investigated the role of actomyosin dynamics in regulating protrusion formation by removing matrix-adhesion sites and disrupting cell contractility.

### Actomyosin dynamics mediate protrusion formation at low critical stresses

To gain insight into the mechanisms involved in hMSC protrusion formation as a function of the nonlinear elastic matrix properties, we first investigated if protrusions were impacted by removing the fibronectin-derived peptide epitope, CRGDS, from the hydrogel. In the PPEGA9-**80** and PPEGA9-**110** samples which span the BRSR, we observed a striking decrease in the length of protrusions per cell and increase in solidity, although the number of protrusions per cell increased (**Figure 5a-d**). While the gels lacked integrin-binding epitopes, hMSCs were still able to extend many actin-rich filipodia, which we attributed to probing the matrix in search of adhesion sites.

**Figure 5.**
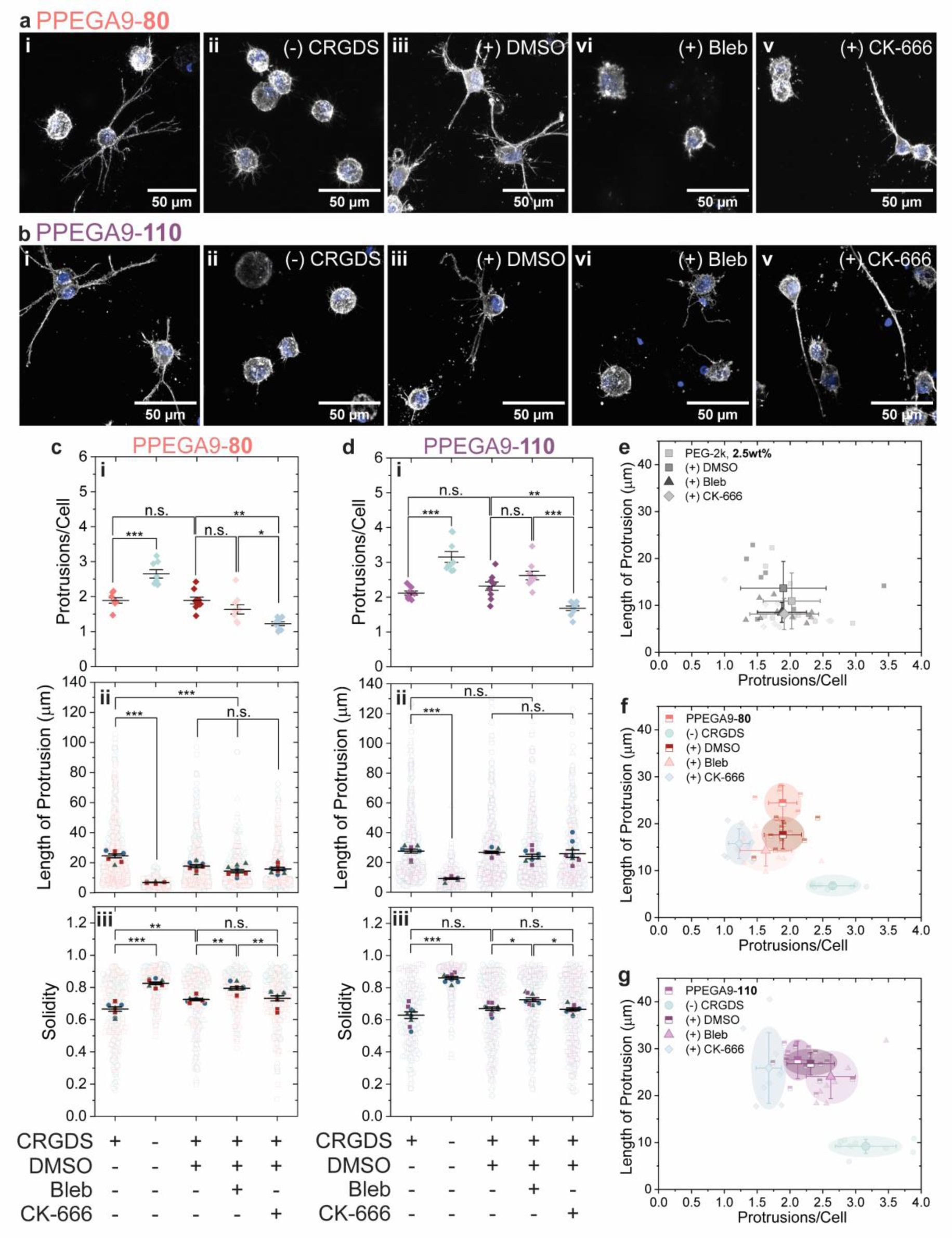
Actomyosin dynamics dictate cellular responses to biologically relevant critical stresses. Representative maximum intensity projections of f-actin (grey) and nuclei (blue) for **a,** PPEGA9-**80** and **b,** PPEGA-**110** cultured in i) growth media, ii) without CRGDS in the network, iii) with DMSO (vehicle), iv) with 50 µM Blebbistatin, and v) with 100 µM CK-666^[38]^ for 3 days of culture in the hydrogels. Image analysis for **c**, PPEGA9-**80** and **d**, PPEGA9-**110** to quantify the changes in **i**) protrusions per cell, **ii**) length of protrusions, and **iii**) solidity for each drug treatment. The number of protrusions per cell plotted with respect to length of protrusions to summarize the shift in morphological features with each treatment for **e,** PEG-2k, **2.5wt%**, **f,** PPEGA9-**80**, and **g,** PPEGA9-**110**. (n = 3 independent biological replicates)

After modifying matrix adhesivity, we inhibited the dramatic actomyosin-based contractility, observed using 3DTFM, in **Figure 4** using blebbistatin. Three days of treatment did not result in statistical differences in the number or length of protrusions per cell compared to the vehicle (**Figure 5c,i-ii**-**d,i-ii**), but the protrusions were notably more filamentous and disorganized compared to those formed in the untreated or vehicle conditions (**Figure 5b,vi**). These differences are reflected in blebbistatin-treated samples by an increase in solidity for both formulations (**Figure 5c,iii-d,iii**), suggesting that contractility plays a role in strain-stiffening mediated actin-organization and protrusion formation. To further disrupt actomyosin dynamics, we blocked Arp2/3, which is known to be involved in actin branching^[38,39]^ and focal adhesion maturation^[40–42]^, using the small molecule inhibitor CK-666. After three days of treatment, protrusion formation was globally suppressed (**Figure 5c,i-d,i**), and rare protrusions that did form existed as single non-branching extensions (**Figure 5a,v-b,v**), in contrast to the dendrite-like protrusions in untreated and vehicle controls (**Figure 5a-b**, **f-g**).

To further explore the complexities of mechanosensing by hMSCs encapsulated in hydrogels with σ_*c*,*equil*_ near the BRSR, we plotted protrusion length versus the number of protrusions per cell and observed how the cell morphotypes clustered as a function of drug treatment (**Figure 5e-g**). In the modulus-matched PEG-2k, **2.5wt%** control (σ_*c*,*equil*_ = 27 Pa), we observed no shifts in the cell population across all treatments (**Figure 5e**, **Figure S19**). This contrasts with the distinct shifts in cell populations observed for hMSCs cultured within the PPEGA9-**80** and PPEGA9-**110** hydrogels (**Figure 5f-g**). In these gels, removal of CRGDS dramatically decreased the length of protrusions but increased the number of protrusions per cell (green cluster). In addition, treatment with CK-666 did not greatly affect the length of protrusions, but instead shifted the population to fewer protrusions per cell (blue cluster). The suppression of actin branching mediated by Arp2/3 stunted nucleation of protrusions and suppressed dendritic branching along those that did form. Blebbistatin treatment did not shift cell morphotype dramatically from the control, but we observed increased solidity for both conditions compared to the vehicle treatment, indicating disrupted protrusion extension and maturation (**Figure 5c,iii-d,iii**). These changes were only observed in hydrogels with σ_*c*,*equil*_ within the BRSR (**Figure S19**). Taken together, we conclude that intracellular actomyosin dynamics play a critical role in cellular responses to nonlinear elastic mechanics, which manifest as the formation of distinct protrusions in these strain-stiffening matrices.

## CONCLUSION

Nonlinear elasticity is an inherent property of the ECM *in vivo,* but its implications in cell mechanotransduction and behavior has been relatively understudied, limited by the scarcity of synthetic materials that recapitulate this biomechanical feedback mechanism and support 3D cell culture. Through the design of tunable materials that isolate cell-dictated strain-stiffening, one can glean insights into the cellular mechanisms that are guided by this fundamental ECM property. We present a controllable and facile synthetic method to incorporate bottlebrush polymers into hydrogel scaffolds through photoinitiated crosslinking, facilitating access of strain-stiffening mechanical properties within a biologically accessible stress regime. These non-fibrous and nonlinear elastic bottlebrush hydrogels and their tunable properties provide a tool to study how cells respond to nonlinear elastic microenvironments and provide a complementary alternative to natural and synthetic fibrous hydrogels. These results reveal that focal adhesions and actomyosin dynamics regulate cellular sensing of and response to nonlinear elastic mechanical properties which mediate an interesting, protrusion-rich phenotype in hMSCs. This fundamental demonstration of how strain-stiffening engineered microenvironments regulate initiation of cellular protrusion formation has great potential for translation to known cell types which contain dendritic processes, such as astrocytes, oligodendrocytes, microglia, pericytes, and osteocytes, in the future. In addition, the systematic incorporation of degradable and stress-relaxing moieties into the strain-stiffening bottlebrush matrix could enable future controlled investigation and improved modeling of nonlinear viscoelastic ECM microenvironments.

## MATERIALS AND METHODS

### Synthetic Materials

2-(Dodecylthiocarbonothioylthio)-2-methylpropionic acid (DDMAP, Sigma-Aldrich), ethylene glycol (Sigma-Aldrich), 4-dimethylaminopyridine (DMAP, Sigma-Aldrich), *N,N’*-diisopropylcarbodiimide (DIC, Sigma), poly(ethylene glycol) methyl ether acrylate [average Mn = 480 Da] (PEGA9, Sigma-Aldrich), 2,2’-Azobis(2-methylpropionitrile) (AIBN, Sigma-Aldrich), N,N-dimethylformamide (DMF, anhydrous, 99.8%, Alfa Aesar), RC Dialysis tubing (3.5 kDa or 8 kDa molecular weight cut-off, Spectrum Spectra/Por 6, Fisher Scientific), HCl (2M, Sigma-Aldrich), tetrahydrofuran (THF, Sigma-Aldrich), methanol (Sigma-Aldrich), Tris(2-carboxyethyl)phosphine hydrochloride (TCEP, Sigma-Aldrich), n-butylamine (99.5%, Sigma-Aldrich), Endo-exo-5-norbornene-2-carboxylic acid (NB-COOH, Sigma-Aldrich), 20 kDa 8-arm poly(ethylene glycol) amine (tripentaerythritol), HCl salt (JenKem, ≥95% substitution), O-(7-Azabenzotriazol-1-yl)-N,N,N’,N’-tetramethyluronium hexafluorophosphate (HATU, ≥99.5% (HPLC), Chem-Impex), *N,N*-Diisopropylethylamine (DIPEA, Sigma-Aldrich), DPBS, no calcium, no magnesium (PBS, Fisher Scientific), Lithium phenyl-2,4,6-trimethylbenzoylphosphinate (LAP, Sigma-Aldrich), CRGDS (Bachem).

### di-DDMAT Synthesis

2-(Dodecylthiocarbonothioylthio)-2-methylpropionic acid (2 eq.; 5.5 mmol) was combined with ethylene glycol (1 eq.; 2.75 mmol), and 4-dimethylaminopyridine (0.2 eq.; 0.55 mmol) were dissolved in DCM (0.4 M) in an oven dried round bottom flask. The reaction mixture was purged with N2 for 10 minutes before adding *N,N’-*diisopropylcarbodiimide (2.4 eq.; 6.6 mmol) and stirring the reaction mixture at room temperature overnight. After stirring overnight, the insoluble urea biproduct was filtered off and the reaction mixture was concentrated using rotary evaporation to yield a dark yellow oil. The di-DDMAT was purified via a gradient silica column with 95:5 to 60:40 hexanes:ethyl acetate as the eluent. The product was isolated as a deep yellow oil and dried under vacuum at room temperature (yield: 45%).

^1^H NMR (400 MHz, CDCl3): δ 4.30 (s, 4H), 3.26 (t, *J =* 7.5 Hz, 4H), 1.68 (s, 12H), 1.66 (m, 4H), 1.38 (p, *J =* 6.2 Hz, 4H), 1.25 (s, 32H), 0.88 (t, *J =* 6.7 Hz, 6H).

^13^C NMR (400 MHz, CDCl3): δ 172.79, 63.34, 55.81, 37.03, 31.92, 29.65, 29.64, 29.57, 29.47, 29.35, 29.12, 29.00, 27.84, 25.27, 22.70.

### SH-PPEGA9-N_bb_-SH Synthesis

Each PPEGA9 bottlebrush was synthesized via reversible addition fragmentation chain transfer polymerization. Poly(ethylene glycol) methyl ether acrylate [average *M*n = 480 Da] (20, 100, 400, or 600 eq.), di-DDMAT (1 eq.), and AIBN (0.1 eq.) was dissolved in anhydrous DMF (1.0 M) in a dry 20 mL scintillation vial capped with a septum. Each reaction mixture was purged with N2 for 15 min prior to heating the reaction mixture at 70 °C overnight. The reaction mixture was quenched in LN2 prior to being exposed to air. The C12H25-PPEGA9-X-C12H25 were dialyzed in methanol (3.5 kDa or 8 kDa MW cut-off RC dialysis tubing), concentrated via rotary evaporation, and dried under high vac overnight yielding viscous yellow oils. The resulting polymers were characterized by ^1^H NMR in CDCl3 and SEC (Agilent Infinity 1200 HPLC, Viscotech I-MBMMW3078 columns, Wyatt T-rEX differential refractive index and Whyatt HELEOS-II MALS detectors) in DMF with 0.05M LiBr.

The trithiocarbonate end-groups were cleaved by aminolysis. The C12H25-PPEGA9-X-C12H25 (2 eq.) was dissolved in THF (0.2 M) with TCEP (1 eq.) and purged with N2 for 5 min. n-butylamine (100 eq.) was added dropwise to each reaction mixture and stirred at room temperature for 2 hours. The yellow color of the reaction mixture was gone after 30 minutes. After 2 hours, the reaction mixtures were moved to dialysis in methanol (3.5 kDa or 8 kDa MW cut-off RC dialysis tubing). After 12 hours, the dialysis media was changed to acidified methanol (pH = 5-3) for one media change to protonate free thiols and discourage disulfide formation. After at least four more media changes, the purified SH-PPEGA9-X-SH were concentrated via rotary evaporation and dried under high vac overnight yielding clear viscous liquids. The trithiocarbonate end-groups were confirmed to be removed with ^1^H NMR in CDCl3. Procedure adapted from Ohnsorg, M. L. *et al.*^[43]^

### 8-arm 20kDa PEG-NB Synthesis

Endo-exo-5-norbornene-2-carboxylic acid (16 eq.; 1.6 mmol), 8-arm 20 kDa PEG-NH2 (1 eq.; 0.1 mmol), and HATU (16 eq.; 1.6 mmol) was dissolved in DMF (0.4 M) in a flame-dried round bottom flask and purged with N2 for 20 minutes before addition the DIPEA (32 eq.; 3.2 mmol) under N2. The reaction mixture was stirred for 24 hours under an N2 atmosphere. After stirring, the reaction mixture was precipitated into cold ethyl ether and isolated by centrifugation (1.5 – 3.0 min at 4.4 rpm). The isolated precipitate was dissolved in methanol and dialyzed progressively against 50:50 methanol:water to 0:100 methanol:water in a 3.5 kDa MW cut-off RC dialysis bag. The purified macromer was isolated via lyophilization and >95% functionalization was quantified via ^1^H NMR in CDCl3. (off-white solid, yield: 92%)

### Sample Gelation Procedure

Stock solutions of SH-PPEGA9-X-SH (30 wt%; 43.7, 17.5, 10.7, or 7.7 mM, for PPEGA9-20, -50, -80, or -110, respectively) or SH-PEG-2k-SH (214 mM, 30 wt%), 20 kDa 8-arm PEG-NB (21.4 mM, 30 wt%), and LAP (2 wt%, 68 mM) were prepared in PBS and mixed and diluted to the appropriate formulations outlined in Supplementary Table S2. 1 mM CRGDS was added to each formulation during cell encapsulation. The samples were then cast in molds on thiolated coverslips (cell encapsulation), between glass slides with 1 mm spacer treated with Rain-X (FRAP), or in a 1 mL syringe barrel (swollen rheology). The solution was irradiated with λ = 365 nm, *I* **=** 5 mW/cm^2^ light for 5 minutes to crosslink the hydrogels. Following photoinitiated crosslinking the gels were swelled in PBS or cell-laden gels were incubated in media.

### Oscillatory Shear Rheology

Rheological measurements were collected using a TA Instruments Ares DHR-3 rheometer equipped with an 8 mm parallel plate geometry (either smooth or sandblasted) and either a quartz plate curing stage equipped UV light guide accessory attached to an Omnicure 1000 light source (λ = 365 nm, *I* **=** 5 mW/cm^2^) or a Peltier plate. Each measurement was performed in triplicate and averaged with the standard deviation shown as the error.

### In Situ Gelation

The storage moduli for each gel formulation were determined *in situ* by loading the hydrogel solution onto the quartz plate curing stage with the smooth 8 mm parallel plate geometry. The storage and loss moduli (*G’, G”*) were measured using an oscillatory strain with an amplitude of 1% and frequency of 1 rad/s. Moduli were measured for 30 seconds before the light source was turned on for 5 minutes and the sample was measure for another 2 minutes following the light source turning off.

### Strain Sweep

Following *in situ* gelation, we performed an oscillatory strain sweep (0.1 – 200%) with a constant frequency of 1 rad/s.

### Frequency Sweep

Following *in situ* gelation, we performed an amplitude sweep (0.1 – 100 rad/s) with a constant 1% strain.

### Swollen Rheology

Each hydrogel formulation was polymerized under 5 mW/cm2 365 nm light (30 uL gels) and swelled in PBS for 24 hours before measuring to allow the gel to reach equilibrium swelling. The storage modulus was measured on a Pelteier plate at 25 °C, with a 8 mm sandblasted parallel plate to avoid slipping. The axial force applied to each sample was kept consistent at 0.5 N and the storage and loss moduli were measured for 180 seconds at 1% strain and frequency of 1 rad/s. The measured storage moduli measurements were averaged (n=3 individual gels) and reported as the equilibrium swollen modulus.

### Fluorescence Recovery After Photobleaching

To estimate the diffusion coefficient for proteins within each hydrogel formulation, fluorescence recovery after photobleaching (FRAP) was performed on pre-made gels (10 µL) equilibrated overnight in fluorophore solution (5 µM FITC-BSA). Each hydrogel was prepared following the gelation procedures outlined above. Photobleaching was performed on a Nikon A1R laser scanning confocal microscope equipped with a CFI Plan Apochromat Lambda D 10X objective (NA = 0.45). FRAP measurements were performed 3-5 times on independent samples (n=2) for each condition. Baseline fluorescence intensity (λex = 488 nm) was acquired for 15 seconds (1 frame per second, 1-2% laser power) before photobleaching a circular region with radius = 50 µm for 5 seconds at 100% laser power. After photobleaching, fluorescence intensity was measured in the circular region for two minutes; data was fit to the equation:

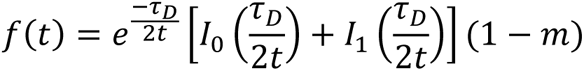

where 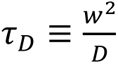. *I*_0_ and *I*_1_ are modified Bessel functions of the first kind, *w* is the radius of the photobleached circular region (50 µm), and (1-*m*) accounts for the steady-state offset. Fitting normalized fluorescence recovery data to this equation using the lsqnonlin solver in MATLAB yields the diffusion coefficient *D* in units of µm^2^/s.

### Biological Materials and Reagents

#### Media Components

Low-glucose (1 ng/mL glucose) Dulbecco’s Modified Eagle Medium (DMEM, ThermoFisher), fetal bovine serum (FBS, ThermoFisher), Penicillin (ThermoFisher), Streptomycin (ThermoFisher) Amphotericin B (ThermoFisher), recombinant human fibroblastic growth factor basic (FGF-2, R&D Systems), dimethyl sulfoxide (anhydrous, Sigma-Aldrich), (-)-Blebbistatin (Bleb, Sigma-Aldrich), CK-666 (≥ 98% (HPLC), Sigma-Aldrich, Lot# 160920)

#### Biological Reagents

Calcein AM (Life Technologies, 0.05 mM), Ethidium Homodimer-1 (Life Technologies, 4.0 mM), Hoechst (1 mg/ml, 1.9 mM), 10% formalin (Sigma-Aldrich), Triton X-100 (Acros), Tween 20 (ThermoFisher), DAPI (Sigma-Aldrich), bovine serum albumin (BSA, Fisher Scientific), Rhodamine Phalloidin (Cytoskeleton, Inc.), anti-YAP antibody IgG2a (mouse, SantaCruz Biotech, sc-101199), recombinant anti-Paxillin antibody [Y113] (rabbit, Abcam, ab32084), goat anti-rabbit AlexaFluor 647 (Invitrogen), goat anti-mouse AlexaFluor 488 (Invitrogen), SPY650-FastAct (Spirochorme), Cytochalasin D (CytoD, Sigma-Aldrich), Tyrode’s Salts (TSS, Sigma-Aldrich)

### hMSC Culture and Encapsulation

The hMSCs were isolated from human bone marrow aspirate (Lonza; donor = 18-year-old black female) and serial expanded following previously described protocols^[44]^. The hMSCs were cultured in low glucose DMEM media supplemented with 10 v/v% FBS, 1 v/v% penicillin-streptomycin, and 0.1 v/v% fungizone with 1.25 ng/mL FGF added after the first day of plating from P2. P3 hMSCs (15 x 10^6^ cells mL^-1^) in media were encapsulated within hydrogel formulations (5 x 10^6^ cells mL^-1^) described above with an additional 1 mM CRGDS. The cell containing hydrogel solution was pipetted into 20 uL molds (5 mm D x 1 mm H) on thiolated glass coverslips. The hydrogels were irradiated with 5 mW/cm^2^ 365 nm light for 5 min before the mold was removed and the cell-laden hydrogel was cultured in 1 mL of media. For the small molecule drug studies, the samples were cultured with 50 µM Blebbistatin, or 100 µM CK-666, following literature precedent,^[38]^ and corresponding vehicle control (DMSO) for 3 days before fixing the samples in 10% formalin.

### Live/Dead Assay

The viability of cells encapsulated within the hydrogels was measured after 1, 3, and 6 days of culture. At the specified timepoints, each cell-laden hydrogel was incubated in 1 mL of 0.5 µM Calcein AM, 4 µM Ethidium Homodimer, and 1.9 µM Hoechst for 20 min at 37 °C. The samples were then washed with PBS and media was replaced before imaging samples on a Nikon Spinning Disk microscope equipped with a CFI Plan Apochromat Lambda D 10X objective (NA = 0.45). Viability was quantified by identifying and counting nuclei in Fiji compared to the number of Calcein AM positive cells vs. Ethidium Homodimer positive nuclei (Supporting information Figure S14-15).

### Immunostaining and Imaging

hMSCs encapsulated in hydrogels were fixed with 10% formalin for 30 minutes followed by three 10-minute washes with PBS. The samples were permeabilized with 0.1% Triton X-100 in PBS for 1 hour at room temperature followed by blocking with 5% BSA for 1 hour at 4 °C. The primary antibody was then added in 5% BSA and incubated overnight at 4 °C [anti-YAP antibody, mouse, (1:250); anti-paxillin antibody, rabbit (1:100)]. The samples were then washed three times with 0.05% Tween 20 in PBS (PBST) for 10 min each. The secondary antibodies were then added in 5% BSA for 1 hour at room temperature [DAPI (1:500), Rhodamine Phalloidin (1:300), goat anti-rabbit 647 (1:500), and goat anti-mouse 488 (IgG3) (1:500)]. The samples were washed three times for 10-minutes with PBST and stored in PBS at 4 °C until imaging.

All samples immunostained for DAPI, Rhodamine Phalloidin, and YAP were imaged using the Perkin Elmer Opera Phenix spinning disc confocal microscope equipped with a 20x water immersion objective (NA = 1) for high throughput quantification.

To obtain higher quality images for subcellular focal adhesion (Paxillin) quantification, samples were also imaged using the Nikon A1R laser scanning confocal microscope equipped with a CFI Apo LWD Lamda S 20XC water immersion objective (NA = 0.95) using Nyquist sampling unless otherwise specified.

### Solidity and Cell Volume Analysis

Cell nuclei were manually segmented and used as markers for marker-controlled watershed segmentation of the cellular bodies in 3D (Supplementary information **Figure S16**). We performed the 3D segmentation in Matlab. We used the Matlab function *regionprops3* to compute the solidity and volume of the segmented cells.

### Blinded Protrusion Analysis

Cell protrusion quantification was performed blinded. In Fiji, maximum intensity projections of the nuclei and actin channels were used to count the number of cells per image and the number of cells with protrusions. The length of each individual protrusion was measured using a line segment tool. The total number of protrusions was then divided by the number of cells with protrusions to calculate the number of protrusions per cell. A total of three images was analyzed per n = 3 biological replicates.

### YAP Analysis

In Fiji, the center most z-plane was identified for each cell before using a line segment drawn across the center of the cell was use generate an intensity vs. location plot for both the YAP and nucleus channels using the *plot profile* tool in Fiji. The nuclear boundaries were defined at the mean half-max of the nuclei intensity. The YAP signal intensity was averaged outside and inside the nucleus and used to calculate the nuclear/cytoplasmic YAP ratio for each sample. Nuclear/cytoplasmic YAP ratios were measured across n = 3 biological replicates, measuring at least 15 cells per replicate.

### Focal Adhesion Quantification

The volume of each focal adhesion per cell was quantified using IMARIS 10.0.0. Each image was analyzed using the ‘Cells’ pathway, to define the nucleus, cell boundary, and “vesicles” (focal adhesions). The focal adhesion volume was quantified using *region growing* tools based on the absolute signal intensity. Each threshold was done manually to ensure accurate focal adhesion quantification for each cell due to the unique morphology with protrusions. The volume of all measured focal adhesions was reported for n = 10 cells.

### Three-Dimensional Traction Force Microscopy

#### Sample Preparation and Data Collection

The contractile behavior of hMSCs within bottlebrush hydrogels were assessed through three-dimensional traction force microscopy (3DTFM)^[45]^. hMSCs were encapsulated in hydrogels following the “hMSC Culture and Encapsulation” protocol outlined above. For 3DTFM, 150,000 cells/mL with 1 v/V% 500 nm fluorescent beads (Floresbrite YG carboxylate beads, Polysciences, Inc.) were encapsulated within each hydrogel on a Nunc glass bottom dish (Nunc Glass, Thermo Scientific) and cultured for 3 days before imaging^[46–48]^. Cells were incubated with SPY650-FastAct (1/300) in media at least 3 hours prior to imaging. Each sample was incubated in Tyrode’s Salt Solution (TSS) with SPY650-FastAct (1/300) for at least 40 min before imaging using a Nikon A1R laser scanning confocal microscope equipped with a CFI Apo LWD Lamda S 20XC water immersion objective (NA = 0.95) and piezo z-drive (z-step size = 0.8 µm, z-stack = 100 µm, 1024 x 1024, zoom = 2.5). The cells were then treated with 4 µM CytoD for 40 minutes and imaged again to collect the relaxed state (Figure S17).

#### Calculation of Bead Displacements

Fluorescent markers were tracked from the CytoD condition to the control condition (TSS) using previously published open-source software^[49]^. In brief, the positions of fluorescent markers were identified, and feature vectors were established for every fluorescent marker which pointed from the centroid of the marker to the five nearest neighboring markers. This process was done for both image sets (CytoD and TSS). Feature vectors serve as a unique identifier for every marker and were compared between the two states to identify matches. This process was repeated until all markers were matched, resulting in the displacement field produced by cellular contractility. The software also corrected for rigid body translations between the image sets. In addition, the software produced surface meshes of the imaged cellular geometries via the marching cubes algorithm^[50]^ which were smoothed in the software Meshlab^[51]^.

The displacement field was interpolated such that local displacements could be ascertained anywhere within the domain of the imaged region. Then, the gradient of the displacement field was computed to obtain the deformation gradient tensor as follows:

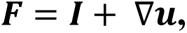

where ***F*** is the deformation gradient tensor, ***I*** is the second-order identity tensor, ***u*** is the displacement field, and ∇***u*** represents the displacement gradient. We considered the hydrogel material to be isotropic and nearly incompressible and modeled their mechanical behavior using a hyperelastic Neo-Hookean model:

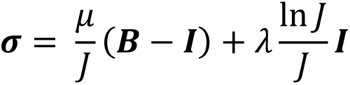

where ***σ*** is the Cauchy stress, *μ* is the first Lamé parameter equivalent to the shear modulus at zero strain, ***B*** is the left Cauchy-Green deformation tensor given by ***B*** = ***FF***^*T*^, *λ* is the second Lamé parameter given by 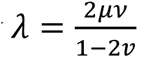 where *v* is Poisson’s ratio (*v* = 0.49), and *J* is the Jacobian given by *J* = det(***F***). The Cauchy stress ***σ*** was computed at each facet comprising the cellular geometry and the resulting traction vector, ***t***, was computed as follows:

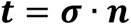

where ***n*** is the outward facing facet normal. Lastly, the total force produced by a cell, *f_tot_*, was calculated using the following equation:

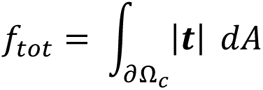

where *∂*Ω_*c*_ is the boundary of the cell and *dA* is an infinitesimal area on *∂*Ω_*c*_.

### Statistical Analysis

Statistical significance was assessed by one-way ANOVA with Tukey’s post hoc. *P>*0.05 was considered to be statistically significant; \**P*<0.05, \*\**P*> 0.01, \*\*\**P>*0.001. Analysis was performed using OriginPro 2022.

## ACKNOWLEDGEMENTS

This work was supported by grants from the National Institutes of Health R01 DE016523 (K.S.A.), T32 AR080630 (M.L.O), and T32 HL007822 (A.K.).

The imaging work was performed at the BioFrontiers Institute Advanced Light Microscopy Core (RRID: SCR_018302). Laser scanning confocal microscopy and FRAP were performed on a Nikon A1R microscope supported by NIST-CU Cooperative Agreement award number 70NANB15H226. We would like to acknowledge and thank the following: Image Analysis Specialist, Jian Wei Tay for his expertise in image analysis using the marker-based watershed segmentation and helpful discussions; Michael Sims at the University of Minnesota for running SEC samples; and Matthew Davidson for helpful expertise and early discussions.

## Table of Contents Figure

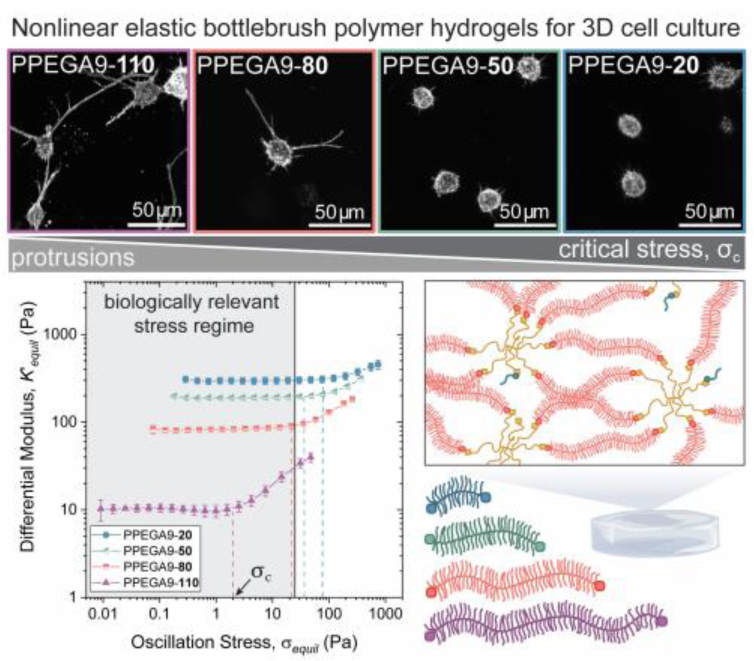

## Table of Contents Text

Nonlinear elastic extracellular matrix properties are difficult to recapitulate without the use of fibrous architectures, which couple strain-stiffening and stress-relaxation. Herein, bottlebrush polymer hydrogels are used to isolate and tune the strain-stiffening biomechanical feedback sensed by human mesenchymal stromal cells (hMSC) in non-fibrous engineered microenvironments. Nonlinear elastic mechanics induced a protrusion-rich hMSC phenotype regulated by focal adhesions and actomyosin dynamics.

## REFERENCES

[1] K. Liu, M. Wiendels, H. Yuan, C. Ruan, P. H. J. Kouwer, Bioact. Mater. 2022, 9, 316.

[2] S. Nam, K. H. Hu, M. J. Butte, O. Chaudhuri, Proc. Natl. Acad. Sci. 2016, 113, 5492.

[3] D. T. Wu, N. Jeffreys, M. Diba, D. J. Mooney, Tissue Eng. Part C Methods 2022, 28, 289.

[4] O. Chaudhuri, J. Cooper-White, P. A. Janmey, D. J. Mooney, V. B. Shenoy, Nature 2020, 584, 535.

[5] C. Storm, J. J. Pastore, F. C. MacKintosh, T. C. Lubensky, P. A. Janmey, Nature 2005, 435, 191.

[6] A. V. Dobrynin, J.-M. Y. Carrillo, Macromolecules 2011, 44, 140.

[7] W. F. M. Daniel, J. Burdyńska, M. Vatankhah-Varnoosfaderani, K. Matyjaszewski, J. Paturej, M. Rubinstein, A. V. Dobrynin, S. S. Sheiko, Nat. Mater. 2016, 15, 183.

[8] R. K. Das, V. Gocheva, R. Hammink, O. F. Zouani, A. E. Rowan, Nat. Mater. 2016, 15, 318.

[9] L.-H. Cai, T. E. Kodger, R. E. Guerra, A. F. Pegoraro, M. Rubinstein, D. A. Weitz, Adv. Mater. 2015, 27, 5132.

[10] S. S. Sheiko, A. V. Dobrynin, Macromolecules 2019, 52, 7531.

[11] A. N. Keith, M. Vatankhah-Varnosfaderani, C. Clair, F. Fahimipour, E. Dashtimoghadam, A. Lallam, M. Sztucki, D. A. Ivanov, H. Liang, A. V. Dobrynin, S. S. Sheiko, ACS Cent. Sci. 2020, 6, 413.

[12] V. G. Reynolds, S. Mukherjee, R. Xie, A. E. Levi, A. Atassi, T. Uchiyama, H. Wang, M. L. Chabinyc, C. M. Bates, Mater. Horiz. 2020, 7, 181.

[13] X. Yu, Y. Wang, M. Li, Y. Zhang, Y. Huang, Q. Qian, Y. Zheng, Q. Hou, X. Fan, ACS Appl. Polym. Mater. 2023, 5, 2750.

[14] K. J. Arrington, S. C. Radzinski, K. J. Drummey, T. E. Long, J. B. Matson, ACS Appl. Mater. Interfaces 2018, 10, 26662.

[15] Y. Tan, H. Chen, W. Kang, X. Wang, Macromolecules 2022, 55, 9715.

[16] M. R. Maw, A. K. Tanas, E. Dashtimoghadam, E. A. Nikitina, D. A. Ivanov, A. V. Dobrynin, M. Vatankhah-Varnosfaderani, S. S. Sheiko, ACS Appl. Mater. Interfaces 2023, 15, 41870.

[17] M. Vatankhah-Varnosfaderani, W. F. M. Daniel, M. H. Everhart, A. A. Pandya, H. Liang, K. Matyjaszewski, A. V. Dobrynin, S. S. Sheiko, Nature 2017, 549, 497.

[18] H. Liang, Z. Wang, A. V. Dobrynin, Macromolecules 2019, 52, 8617.

[19] M. Vatankhah-Varnosfaderani, A. N. Keith, Y. Cong, H. Liang, M. Rosenthal, M. Sztucki, C. Clair, S. Magonov, D. A. Ivanov, A. V. Dobrynin, S. S. Sheiko, Science 2018, 359, 1509.

[20] F. Vashahi, M. R. Martinez, E. Dashtimoghadam, F. Fahimipour, A. N. Keith, E. A. Bersenev, D. A. Ivanov, E. B. Zhulina, P. Popryadukhin, K. Matyjaszewski, M. Vatankhah-Varnosfaderani, S. S. Sheiko, Sci. Adv. 2022, 8, eabm2469.

[21] P. H. J. Kouwer, M. Koepf, V. A. A. Le Sage, M. Jaspers, A. M. van Buul, Z. H. Eksteen-Akeroyd, T. Woltinge, E. Schwartz, H. J. Kitto, R. Hoogenboom, S. J. Picken, R. J. M. Nolte, E. Mendes, A. E. Rowan, Nature 2013, 493, 651.

[22] M. Fernández-Castaño Romera, R. Göstl, H. Shaikh, G. Ter Huurne, J. Schill, I. K. Voets, C. Storm, R. P. Sijbesma, J. Am. Chem. Soc. 2019, 141, 1989.

[23] E. Prince, E. Kumacheva, Nat. Rev. Mater. 2019, 4, 99.

[24] M. D. Davidson, M. E. Prendergast, E. Ban, K. L. Xu, G. Mickel, P. Mensah, A. Dhand, P. A. Janmey, V. B. Shenoy, J. A. Burdick, Sci. Adv. 2021, 7, eabi8157.

[25] M. Jaspers, M. Dennison, M. F. J. Mabesoone, F. C. MacKintosh, A. E. Rowan, P. H. J. Kouwer, Nat. Commun. 2014, 5, 5808.

[26] K. Liu, S. M. Mihaila, A. Rowan, E. Oosterwijk, P. H. J. Kouwer, Biomacromolecules 2019, 20, 826.

[27] P. Dontula, C. W. Macosko, L. E. Scriven, AIChE J. 1998, 44, 1247.

[28] I. N. Haugan, M. J. Maher, A. B. Chang, T.-P. Lin, R. H. Grubbs, M. A. Hillmyer, F. S. Bates, ACS Macro Lett. 2018, 7, 525.

[29] V. Satopää, J. Albrecht, D. Irwin, B. Raghavan, in 2011 31st Int. Conf. Distrib. Comput. Syst. Workshop, IEEE, Minneapolis, MN, USA, 2011, pp. 166–171.

[30] R. C. Ollier, Y. Xiang, A. M. Yacovelli, M. J. Webber, Chem. Sci. 2023, 14, 4796.

[31] E. Prince, S. Morozova, Z. Chen, V. Adibnia, I. Yakavets, S. Panyukov, M. Rubinstein, E. Kumacheva, Proc. Natl. Acad. Sci. 2023, 120, e2220755120.

[32] M. W. Tibbitt, K. S. Anseth, Biotechnol. Bioeng. 2009, 103, 655.

[33] H. Yuan, K. Liu, M. Cóndor, J. Barrasa-Fano, B. Louis, J. Vandaele, P. de Almeida, Q. Coucke, W. Chen, E. Oosterwijk, C. Xing, H. Van Oosterwyck, P. H. J. Kouwer, S. Rocha, Proc. Natl. Acad. Sci. 2023, 120, e2216934120.

[34] J. Kumari, R. Hammink, J. Baaij, F. A. D. T. G. Wagener, P. H. J. Kouwer, Biomater. Adv. 2024, 156, 213705.

[35] S. Natan, Y. Koren, O. Shelah, S. Goren, A. Lesman, Mol. Biol. Cell 2020, 31, 1474.

[36] S. Dupont, L. Morsut, M. Aragona, E. Enzo, S. Giulitti, M. Cordenonsi, F. Zanconato, J. Le Digabel, M. Forcato, S. Bicciato, N. Elvassore, S. Piccolo, Nature 2011, 474, 179.

[37] C. Yang, M. W. Tibbitt, L. Basta, K. S. Anseth, Nat. Mater. 2014, 13, 645.

[38] K. Bera, A. Kiepas, I. Godet, Y. Li, P. Mehta, B. Ifemembi, C. D. Paul, A. Sen, S. A. Serra, K. Stoletov, J. Tao, G. Shatkin, S. J. Lee, Y. Zhang, A. Boen, P. Mistriotis, D. M. Gilkes, J. D. Lewis, C.-M. Fan, A. P. Feinberg, M. A. Valverde, S. X. Sun, K. Konstantopoulos, Nature 2022, 611, 365.

[39] A. Giri, S. Bajpai, N. Trenton, H. Jayatilaka, G. D. Longmore, D. Wirtz, FASEB J. 2013, 27, 4089.

[40] D. S. Chorev, O. Moscovitz, B. Geiger, M. Sharon, Nat. Commun. 2014, 5, 3758.

[41] R. T. Böttcher, M. Veelders, P. Rombaut, J. Faix, M. Theodosiou, T. E. Stradal, K. Rottner, R. Zent, F. Herzog, R. Fässler, J. Cell Biol. 2017, 216, 3785.

[42] R. Zhao, S. Cui, Z. Ge, Y. Zhang, K. Bera, L. Zhu, S. X. Sun, K. Konstantopoulos, Sci. Adv. 2021, 7, eabg4934.

[43] M. L. Ohnsorg, P. C. Prendergast, L. L. Robinson, M. R. Bockman, F. S. Bates, T. M. Reineke, ACS Macro Lett. 2021, 10, 375.

[44] V. V. Rao, M. K. Vu, H. Ma, A. R. Killaars, K. S. Anseth, Bioeng. Transl. Med. 2019, 4, 51.

[45] W. R. Legant, J. S. Miller, B. L. Blakely, D. M. Cohen, G. M. Genin, C. S. Chen, Nat. Methods 2010, 7, 969.

[46] A. Khang, E. Lejeune, A. Abbaspour, D. P. Howsmon, M. S. Sacks, J. Biomech. Eng. 2021, 143, 094503.

[47] A. Khang, Q. Nguyen, X. Feng, D. P. Howsmon, M. S. Sacks, Acta Biomater. 2023, 163, 194.

[48] R. Tuscher, A. Khang, T. M. West, C. Camillo, G. Ferrari, M. S. Sacks, Front. Physiol. 2023, 14, 1168691.

[49] E. Lejeune, A. Khang, J. Sansom, M. S. Sacks, SoftwareX 2020, 11, 100417.

[50] S. Van Der Walt, J. L. Schönberger, J. Nunez-Iglesias, F. Boulogne, J. D. Warner, N. Yager, E. Gouillart, T. Yu, PeerJ 2014, 2, e453.

[51] P. Cignoni, M. Callieri, M. Corsini, M. Dellepiane, F. Ganovelli, G. Ranzuglia, Eurographics Ital. Chapter Conf. 2008, 129.

