## Supplementary Material for "Nonlinear Elastic Bottlebrush Polymer Hydrogels Modulate Actomyosin Mediated Protrusion Formation in Mesenchymal Stromal Cells"

**Supplementary Information Section 1:** Bottlebrush polymer synthesis and characterization

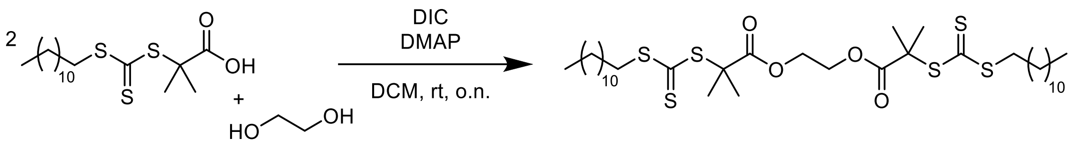


**Scheme S1.** Synthetic scheme for di-DDMAT synthesis

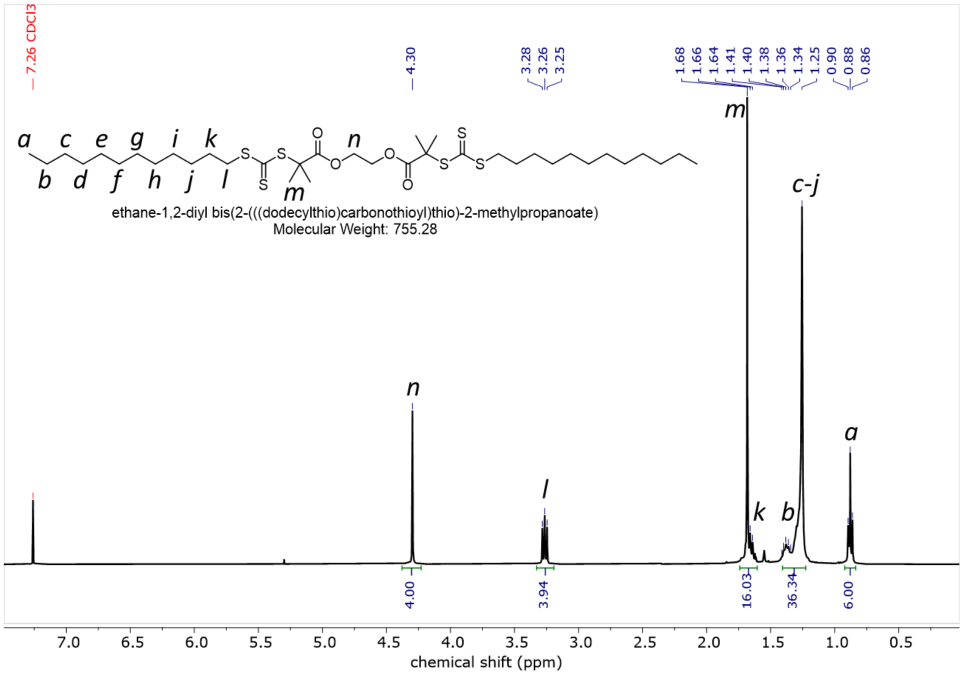


**Figure S1.** $^{1}H$ NMR of di-DDMAT in $CDCl_3$

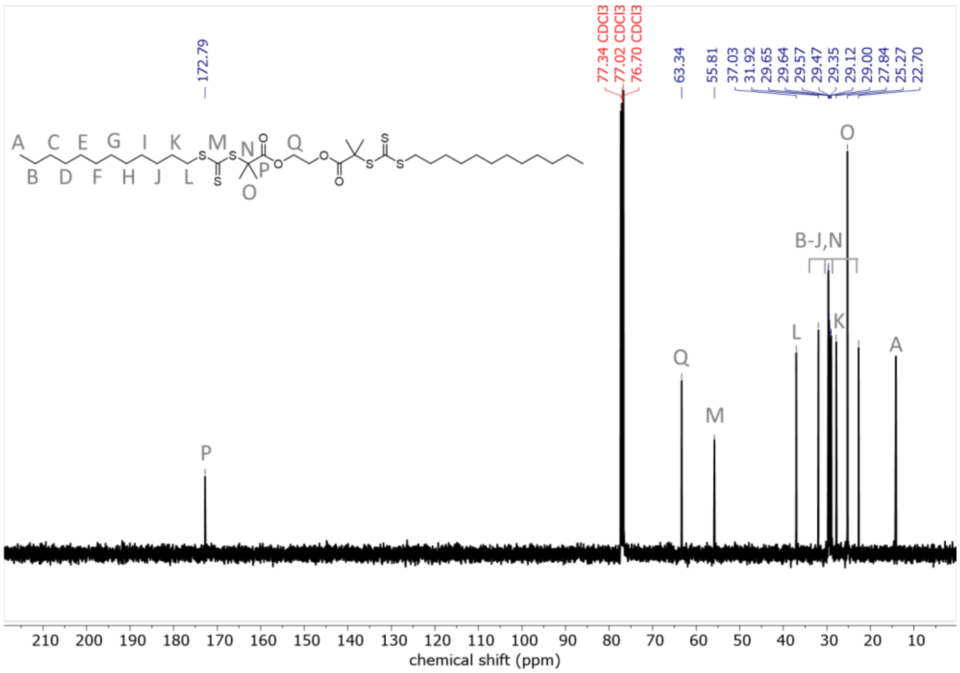


**Figure S2.** $^{13}C$ NMR of di-DDMAT in $CDCl_3$

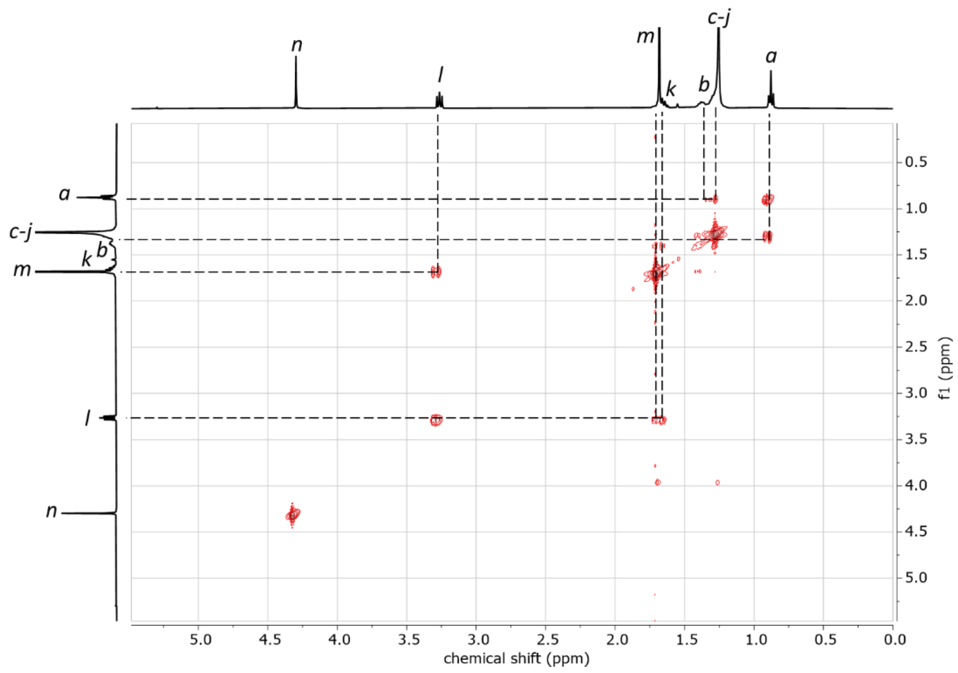


**Figure S3.** COSY NMR of di-DDMAT in $CDCl_3$

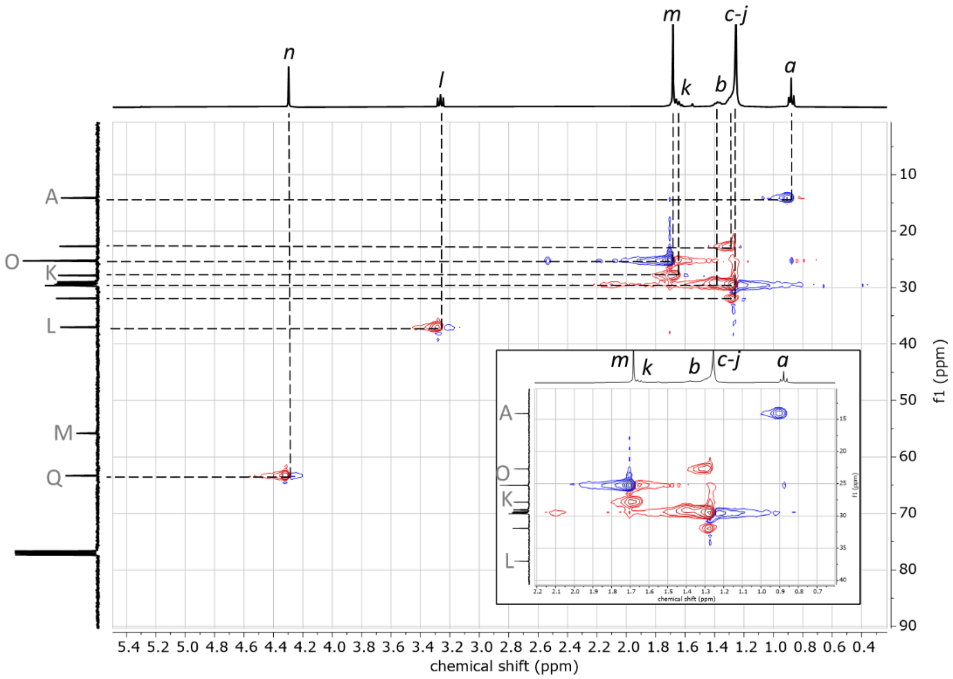


**Figure S4.** HSQC NMR of di-DDMAT in $CDCl_3$

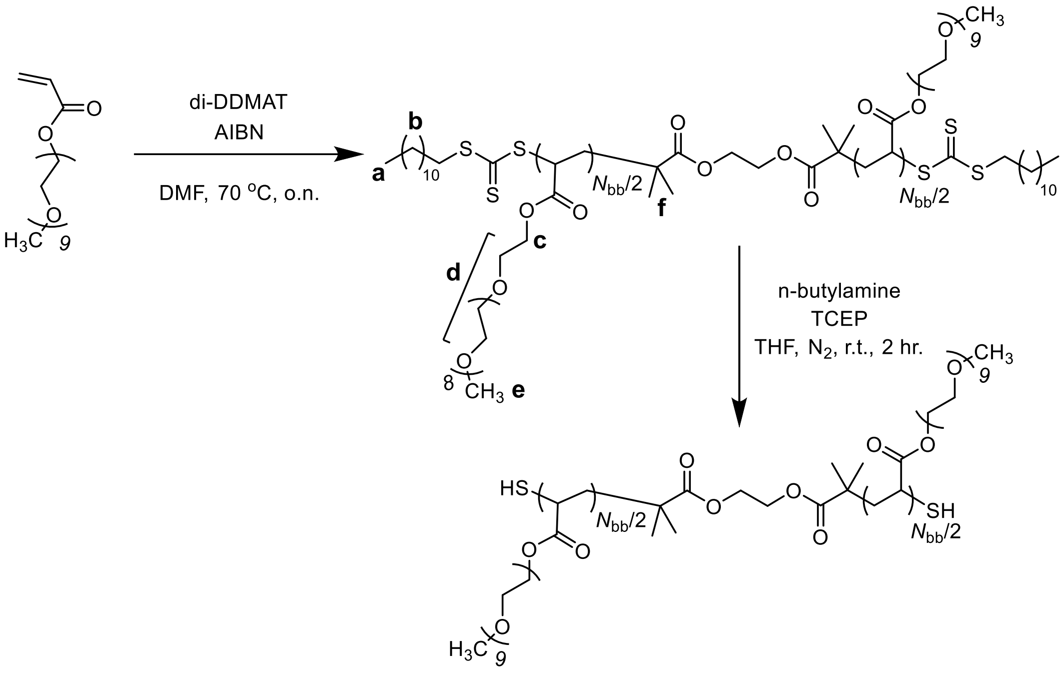


**Scheme S2.** RAFT polymerization of PPEGA9 using di-DDMAT and subsequent aminolysis to yield SH-PPEGA9-$\boldsymbol{N}_{\mathbf{bb}}$-SH

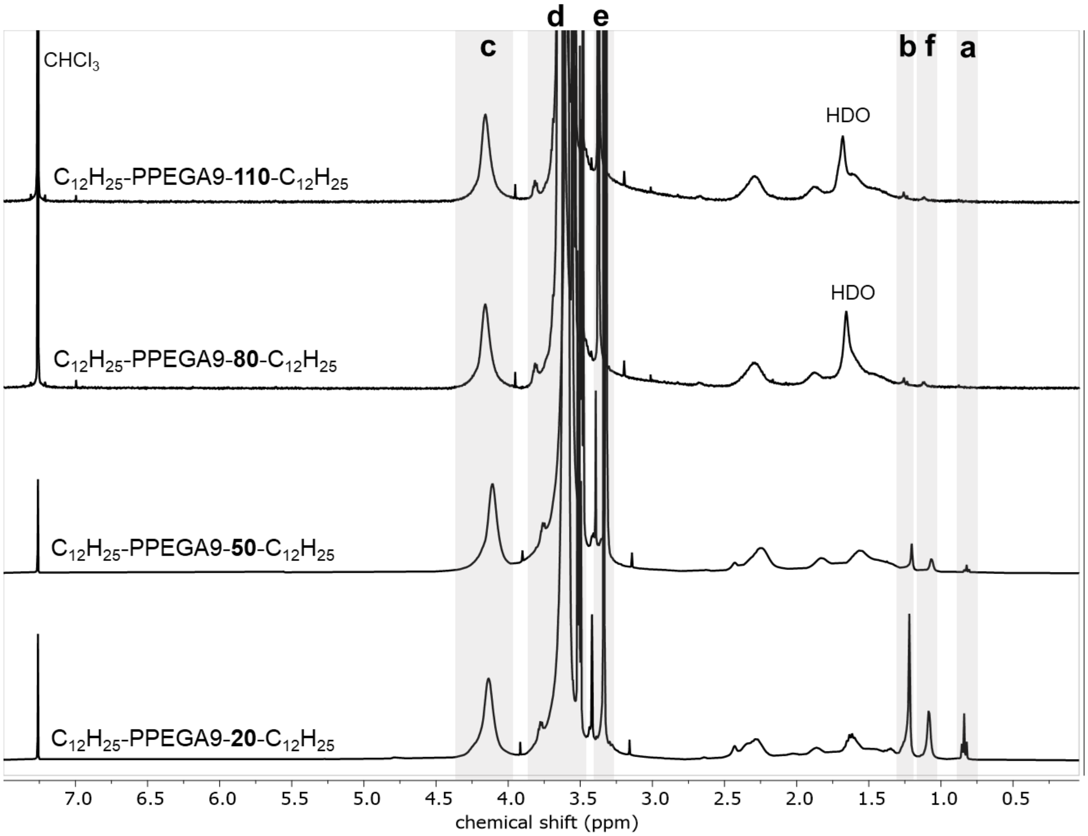


**Figure S5.** $^{1}$H NMR Stack of $C_{12}H_{25}$-PPEGA9-**20-110**-$C_{12}H_{25}$

**Table S1.** Molecular weight summary of all $C_{12}H_{25}$-PPEGA9-X-$C_{12}H_{25}$ bottlebrushes analyzed by SEC in DMF with 0.05M LiBr

| Sample Name | CTA:PEGA9 | $M_w$ (kDa)[a] | $M_n$ (kDa)[a] | Đ[a] | $N_{bb}$ |
|---|---|---|---|---|---|
| $C_{12}H_{25}$-PPEGA9-**20**-$C_{12}H_{25}$ | 1:25 | 11.0 | 9.8 | 1.12 | 19 |
| $C_{12}H_{25}$-PPEGA9-**50**-$C_{12}H_{25}$ | 1:100 | 30.3 | 24.5 | 1.24 | 49 |
| $C_{12}H_{25}$-PPEGA9-**80**-$C_{12}H_{25}$ | 1:400 | 58.6 | 40.2 | 1.46 | 82 |
| $C_{12}H_{25}$-PPEGA9-**110**-$C_{12}H_{25}$ | 1:600 | 104.0 | 55.3 | 1.88 | 114 |

[a)] SEC-MALS in DMF with 0.05M LiBr

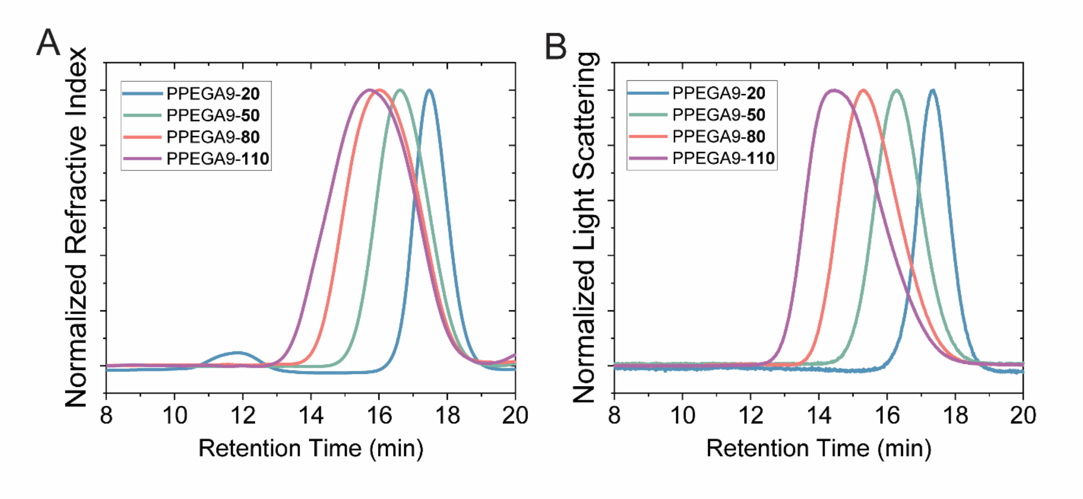


**Figure S6.** SEC traces of $C_{12}H_{25}$-PPEGA9-X-$C_{12}H_{25}$ samples in DMF with 0.05M LiBr. (A) normalized refractive index and (B) normalized light scattering intensities plotted as a function of retention time.

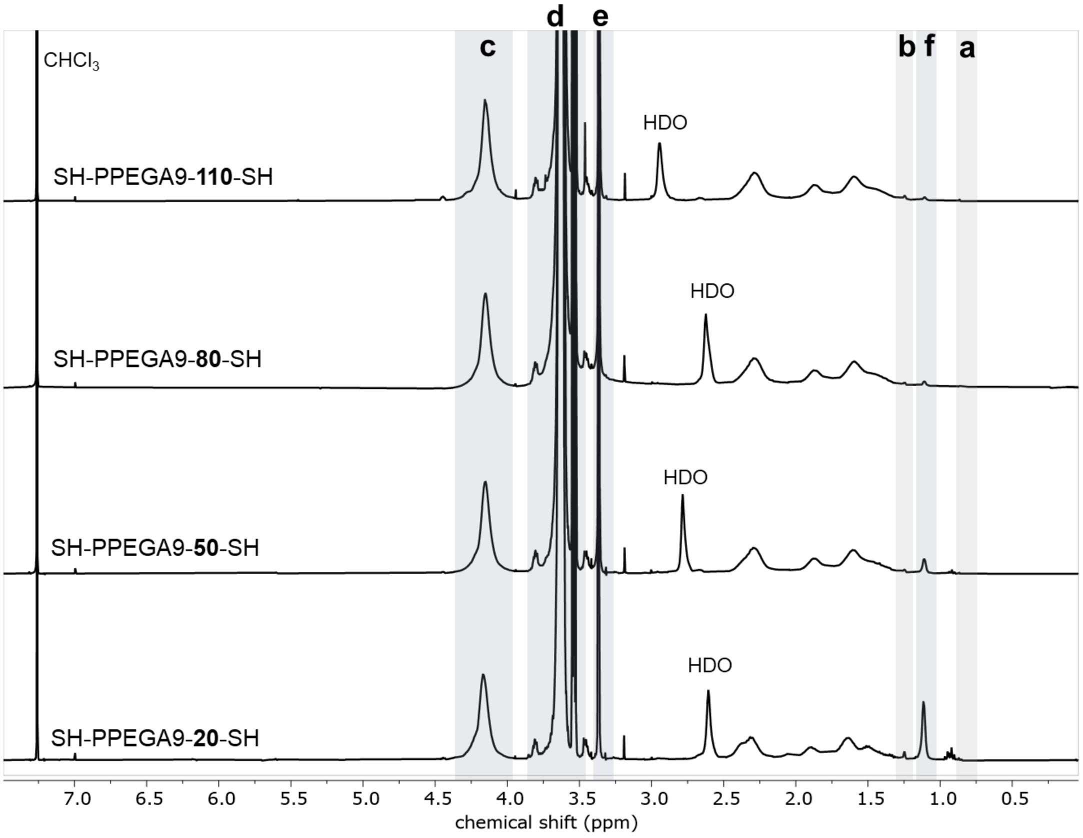


**Figure S7.** $^1$H NMR Stack of SH-PPEGA9-**20-110**-SH

**Supplementary Information Section 2:** Hydrogel formulation and characterization

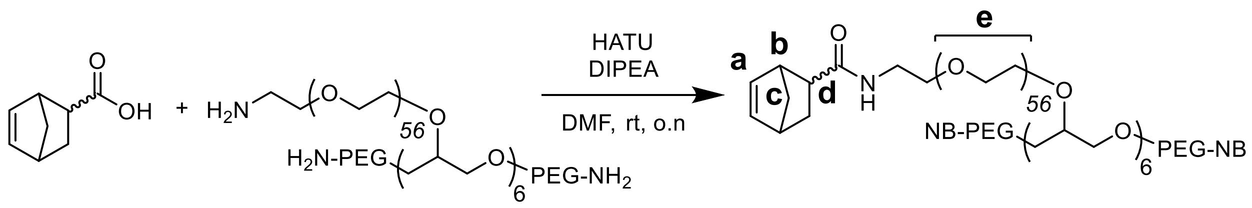


**Scheme S3.** 8-arm 20kDa PEG-NB synthesis

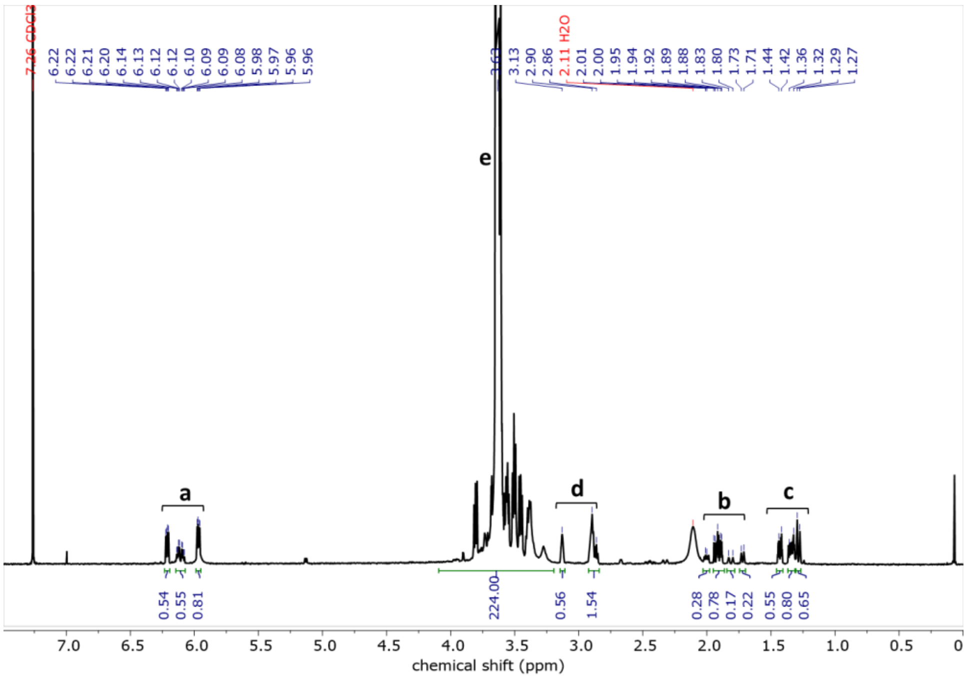


**Figure S8.** $^{1}$H NMR of 8-arm 20kDa PEG-NB

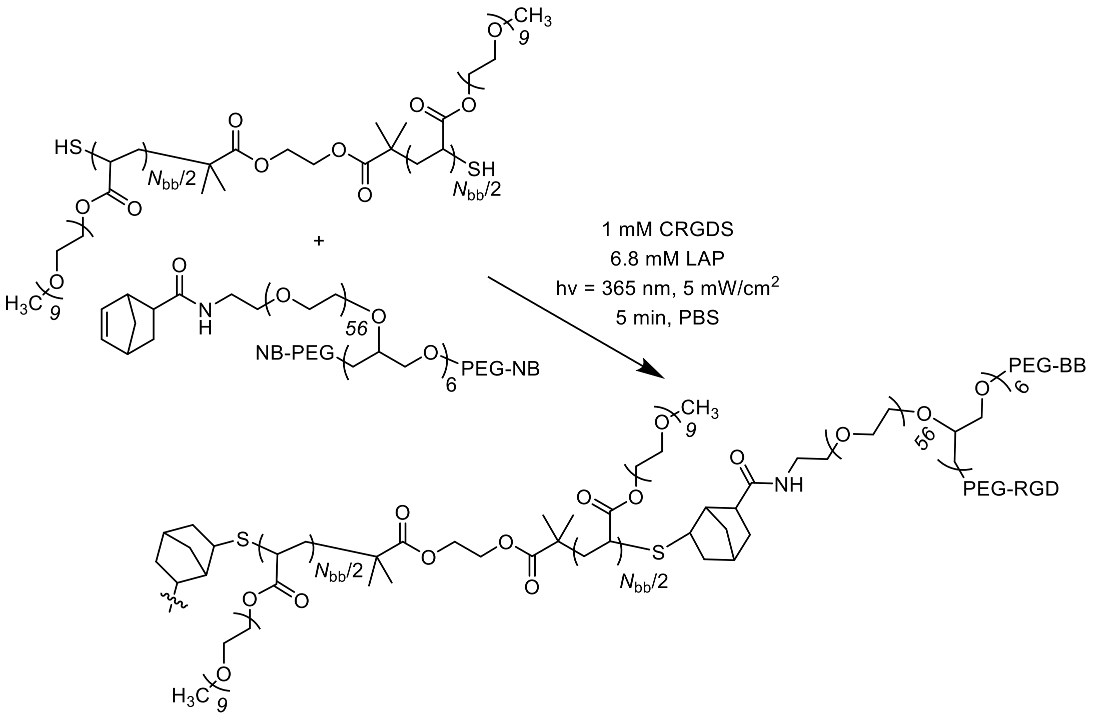


**Scheme S4.** Thiol-ene photo-crosslinking of SH-PPEGA9-$\boldsymbol{N}_{\mathbf{bb}}$-SH and 8-arm 20kDa PEG-NB to form bottlebrush polymer networks.

**Table S2.** Hydrogel formulation summary

| **Dithiol Macromer** | $M_{n, SEC}$[a] [kDa] | SH:NB | Initial wt%[b] | Dithiol conc.[c] [mM] | 8-arm 20kDa PEG-NB conc.[c] [mM] | CRGDS conc.[d] [mM] | LAP conc.[e] [mM] | $G'_{equil}$[f] [Pa] | Diffusion coefficient [um²/s][g] |
|---|---|---|---|---|---|---|---|---|---|
| SH-PPEGA9-**20**-SH | 9.8 | 1.0:1.0 | 10.0 | 9.6 | 2.5 | 1.0 | 6.8 | 311 ± 30 | 13.5 ± 1.3 |
| SH-PEG-2k-SH | 2.0 | 1.0:1.0 | 3.5 | 6.8 | 1.8 | 1.0 | 6.8 | 180 ± 36 | 13.1 ± 1.6 |
| SH-PPEGA9-**50**-SH | 24.5 | 1.0:1.0 | 10.0 | 4.9 | 1.2 | 1.0 | 6.8 | 197 ± 5 | 17.5 ± 0.6 |
| SH-PEG-2k-SH | 2.0 | 1.0:1.0 | 2.5 | 4.9 | 1.3 | 1.0 | 6.8 | 112 ± 11 | 15.3 ± 1.2 |
| SH-PPEGA9-**80**-SH | 40.2 | 1.0:1.0 | 10.0 | 3.2 | 0.7 | 1.0 | 6.8 | 90 ± 4 | 20.6 ± 0.5 |
| SH-PPEGA9-**110**-SH | 55.3 | 1.0:1.2 | 10.5 | 2.5 | 0.7 | 1.0 | 6.8 | 12 ± 3 | 22.6 ± 0.8 |

a) SEC-MALS in DMF with 0.05M LiBr. b) Total weight percent of macromer, SH-PPEGA9-$N_{bb}$-SH or SH-PEG-2k-SH and 8-arm 20kDa PEG-NB, in the formulation upon gelation before swelling. c) Final concentration diluted from 30 wt% stock solutions. d) Final concentration diluted from 70 mM stock solution. e) Final concentration diluted from 68 mM (2 wt%) stock solution. f) Measured by oscillatory shear rheology using a sandblasted 8 mm parallel plate geometry at a constant axial force of 0.5 N. g) Calculated diffusion coefficient for 5 µM FITC-BSA within the equilibrium swollen hydrogel (FRAP).

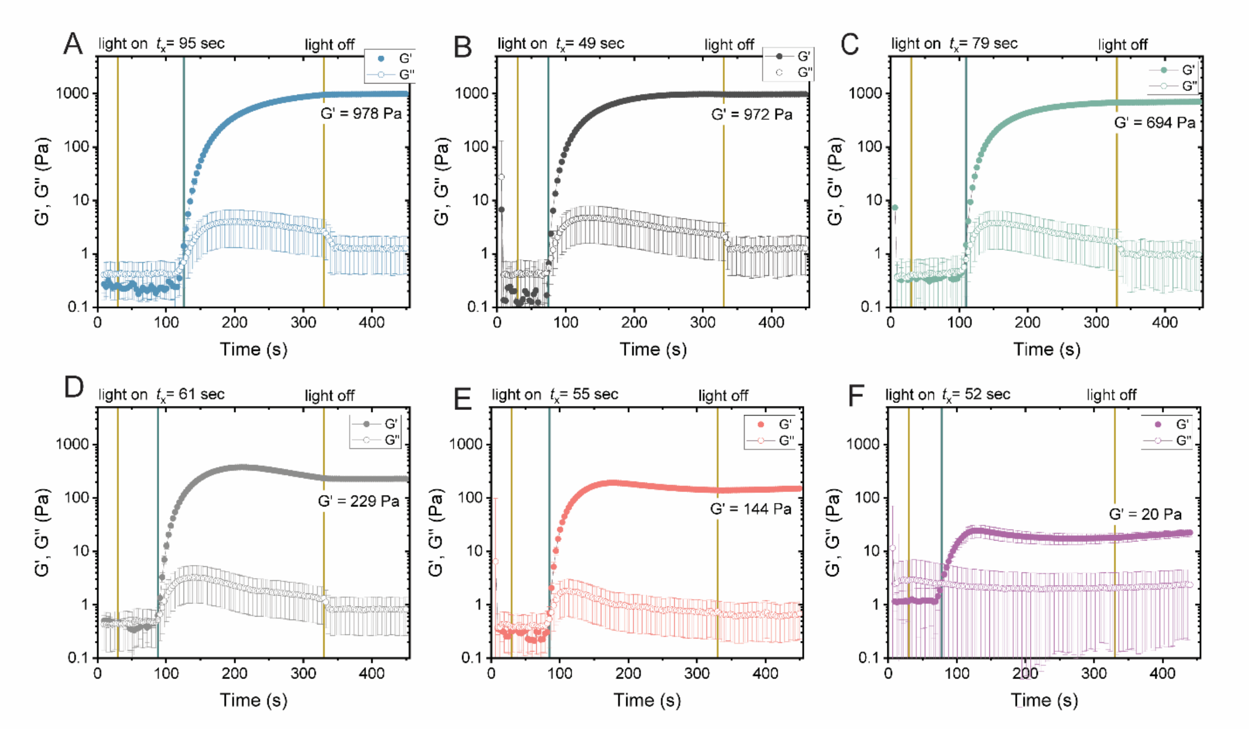


**Figure S9.** *In situ* gelation plots – showing network evolution with time for each hydrogel formulation, (A) PPEGA9-**20**, (B) PEG-2k, **3.5wt%**, (C) PPEGA9-**50**, (D) PEG-2k, **2.5 wt%**, (E) PPEGA9-**80**, and (F) PPEGA9-**110**. $t_x$ designates the crossover time (sec) where *G'* overtakes *G"* after the 365 nm light has been turned on. Data represents the mean ± SD across at least n = 3 replicates.

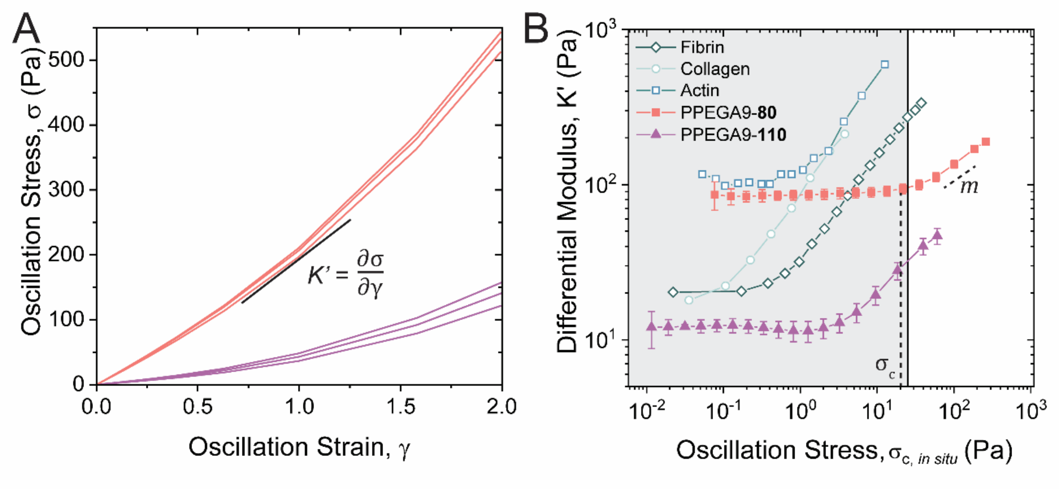


**Figure S10.** (A) oscillatory stress vs oscillatory strain curves for the bottlebrush hydrogels with $\sigma_{c,equil}$ within the BRSR. PPEGA9-**80** (salmon) and PPEGA9-**110** (purple). (B) Overlay of bottlebrush hydrogel formulations which have $\sigma_{c,equil}$ within the BRSR with biological fibrous nonlinear elastic components such as, fibrin, collagen, and actin. Differential modulus vs. oscillation stress data was mined from Das *et al.*[1] from the original data set in Storm *et al.*[2]

*Equilibrium/In situ Scaling:*

The nonlinear elastic properties of the hydrogels were measured directly after network formation (Fig. **S9**) using a strain sweep from 0-200% strain at 1 rad/s (Fig. **S12**). From the resulting oscillation stress versus oscillation strain curves (Fig. **2a** and **S10A**), the derivative yielded the differential modulus, *K'*, and was plotted as a function of oscillation stress (Fig. **S11A**). The critical stress, or onset of network stiffening, ($\sigma_{c,in\ situ}$) was observed so be linearly correlated ($r^2 = 0.98$) to $G'_{in\ situ}$ (Fig. **S11B**). Therefore, we estimated the equilibrium critical stress ($\sigma_{c,equil}$) based on the $G'_{equil}$ using equation 1 (see Table **S3**).

$$\frac{G'_{equil}}{G'_{in\ situ}} * \sigma_{c,in\ situ} = \sigma_{c,equil} \qquad (1)$$

When $\sigma_{c,equil}$ was plotted as a function of $G'_{equil}$ in Fig. **2d** it scales along the same linear correlation as plotted in Fig. **S11B**. *In situ* measurements resulted in only PPEGA9-**110** with a $\sigma_{c,in\ situ}$ within the BRSR, however when scaled based on the $G'_{equil}$, both the PPEGA0-**80** and PPEGA9-**110** samples had $\sigma_{c,equil}$ within the BRSR (Fig. **2**). The slope of the line after the onset of stiffening, stiffening index, *m*, does not scale with G' and indicates that the lower the critical stress, the more stress sensitive the sample is resulting in almost 3x the amount stiffening per Pa of stress applied in the PPEGA9-**110** sample (Fig. **S11C**, Fig. **2e**).

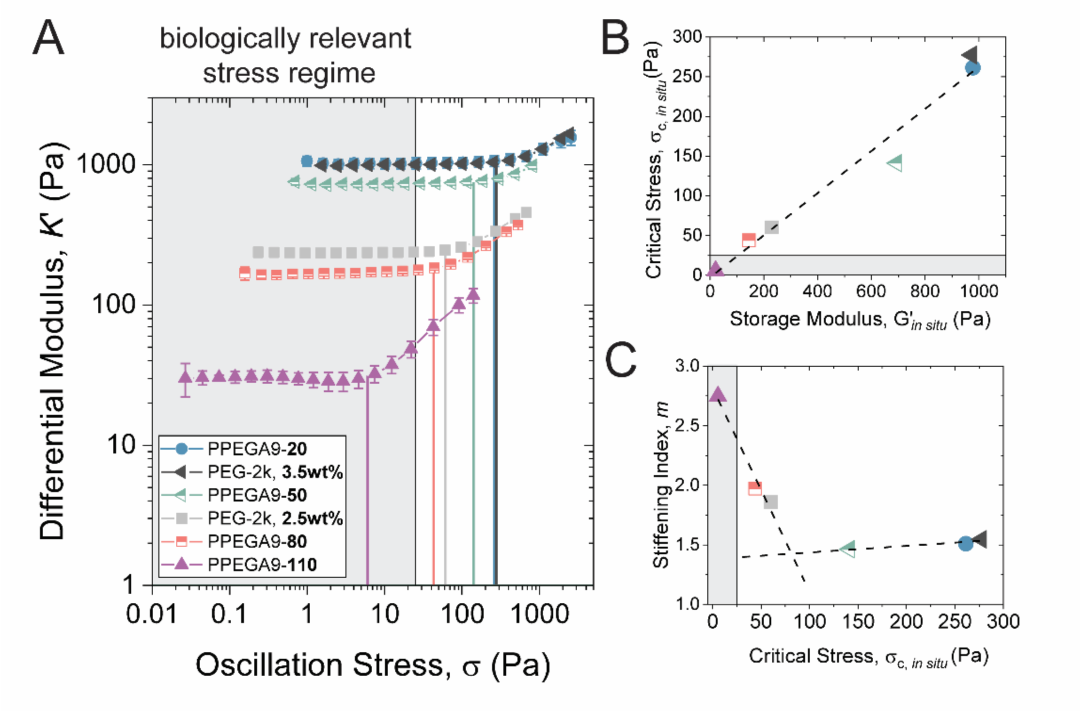


**Figure S11.** *In situ* measured (A) differential modulus, (B) critical stress, and (C) stiffening index plotted against the in *in situ* oscillation stress, storage modulus, and critical stress, respectively. Data represents the mean ± SD across at least n = 3 replicates.

**Table S3.** *Equilibrium/In situ* scaling of hydrogel mechanical properties

| **Dithiol Macromer** | Initial wt%[a] | $G'_{in\ situ}$[b] [Pa] | $\sigma_{c,in\ situ}$[c] [Pa] | *In situ/equil* scaling[d] | $G'_{equil}$[e] [Pa] | $\sigma_{c,equil}$[f] [Pa] | *m* |
|---|---|---|---|---|---|---|---|
| SH-PPEGA9-**20**-SH | 10.0 | 978 ± 82 | 261 ± 23 | 3.446 | 311 ± 30 | 76.0 ± 7.4 | 1.5 ± 0.1 |
| SH-PEG-2k-SH | 3.5 | 972 ± 33 | 277 ± 130 | 5.948 | 180 ± 36 | 49.3 ± 28.6 | 1.5 ± 0.2 |
| SH-PPEGA9-**50**-SH | 10.0 | 694 ± 30 | 141 ± 45 | 3.841 | 197 ± 5 | 36.5 ± 9.7 | 1.5 ± 0.1 |
| SH-PEG-2k-SH | 2.5 | 229 ± 17 | 60 ± 5 | 2.213 | 112 ± 11 | 27.3 ± 2.6 | 1.5 ± 0.01 |
| SH-PPEGA9-**80**-SH | 10.0 | 144 ± 34 | 44 ± 2 | 2.040 | 90 ± 4 | 21.6 ± 0.8 | 2.0 ± 0.02 |
| SH-PPEGA9-**110**-SH | 10.5 | 25 ± 2 | 6 ± 2 | 2.940 | 12 ± 3 | 2.0 ± 0.4 | 2.8 ± 0.1 |

[a)] Total weight percent of macromer, SH-PPEGA9-$N_{bb}$-SH or SH-PEG-2k-SH and 8-arm 20kDa PEG-NB, in the formulation upon gelation before swelling. [b)] Measured by oscillatory shear rheology using an 8 mm parallel plate geometry on a quartz curing stage. [c)] Determined from knee/elbow analysis of K' vs. $\sigma$ plots. [d]$G'_{equil}/G'_{in\ situ}$ from equation 1 [e)] Measured by oscillatory shear rheology using a sandblasted 8 mm parallel plate geometry at a constant axial force of 0.5 N. [f)] Calculated using equation 1.

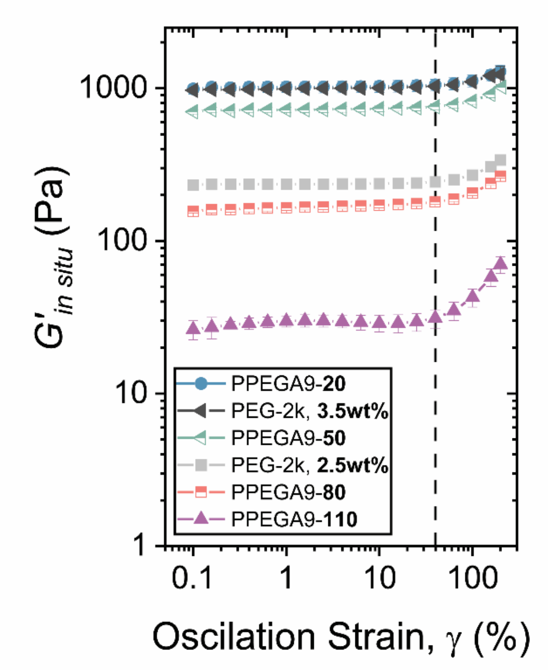


**Figure S12.** Measured $G'_{in\ situ}$ with increasing oscillation strain, %. The dotted line indicates the critical oscillation strain, $\gamma_c$, observed across all samples to be 40%.

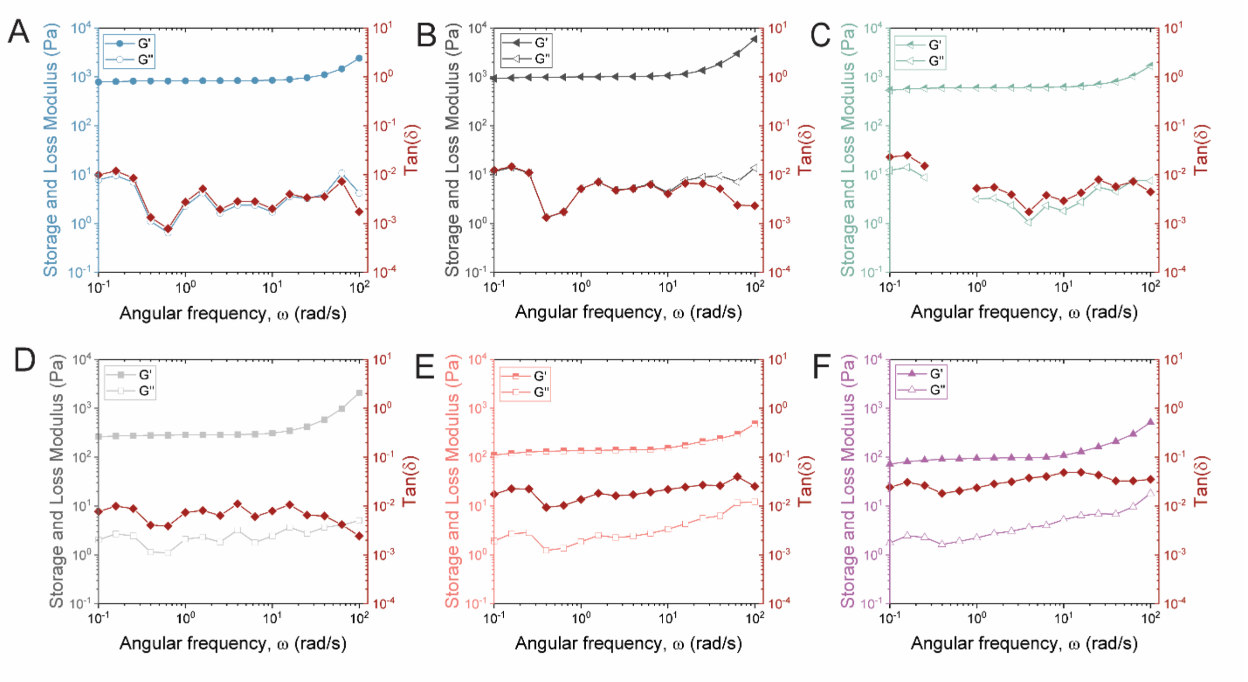


**Figure S13.** Representative frequency sweep (1-100 rad/s) at a constant strain of 1% for each hydrogel formulation (A) PPEGA9-**20**, (B) PEG-2k, **3.5wt%**, (C) PPEGA9-**50**, (D) PEG-2k, **2.5 wt%**, (E) PPEGA9-**80**, and (F) PPEGA9-**110**.

**Supplementary Information Section 3:** Cell culture and image analysis

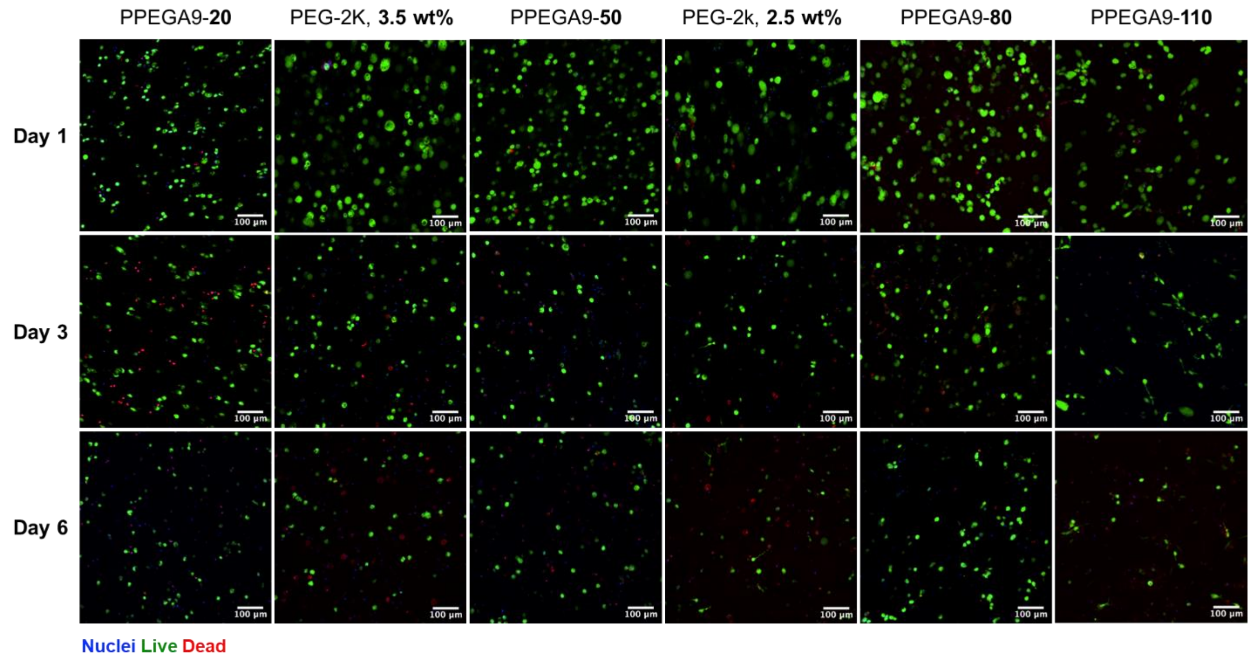


**Figure S14.** Representative confocal images of live/dead assay across all hydrogel formulations on Day 1, 3, and 6. (scale bar = 100 μm)

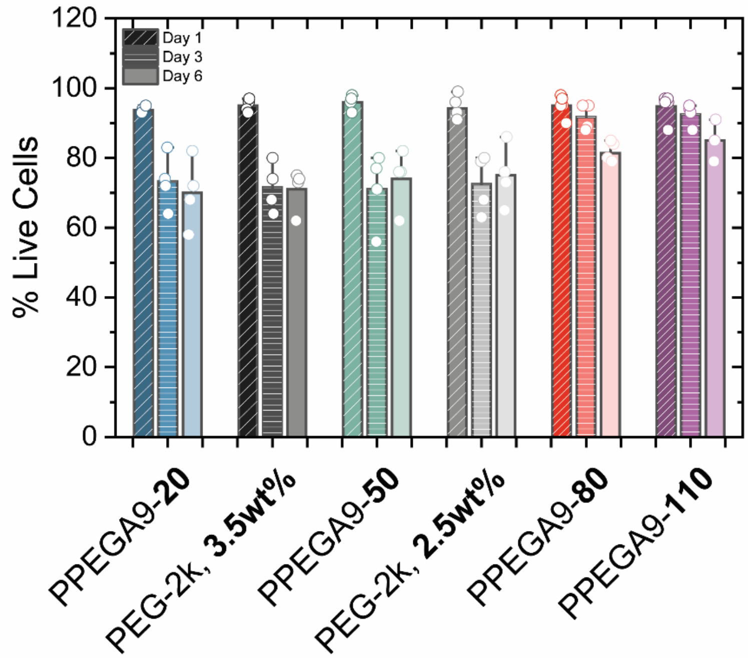


**Figure S15.** Quantified viability for hMSCs encapsulated within each hydrogel condition across days 1, 3, and 6 of culture. (n ≥ 3 imaged regions within each hydrogel)

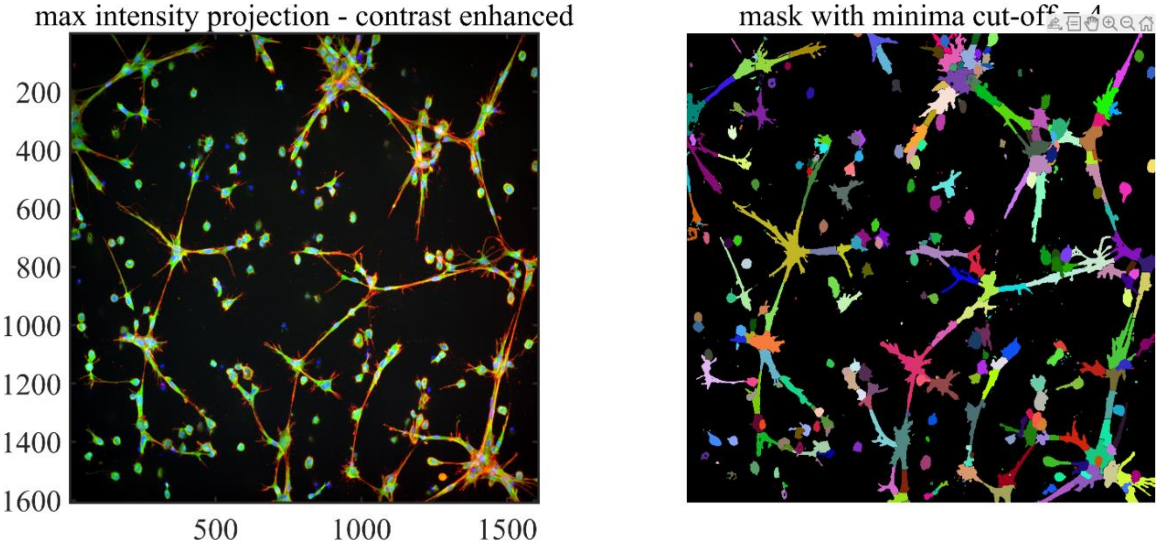


**Figure S16.** Representative marker-base watershed segmentation data showing successful segmentation of a mix of connected and single cells.

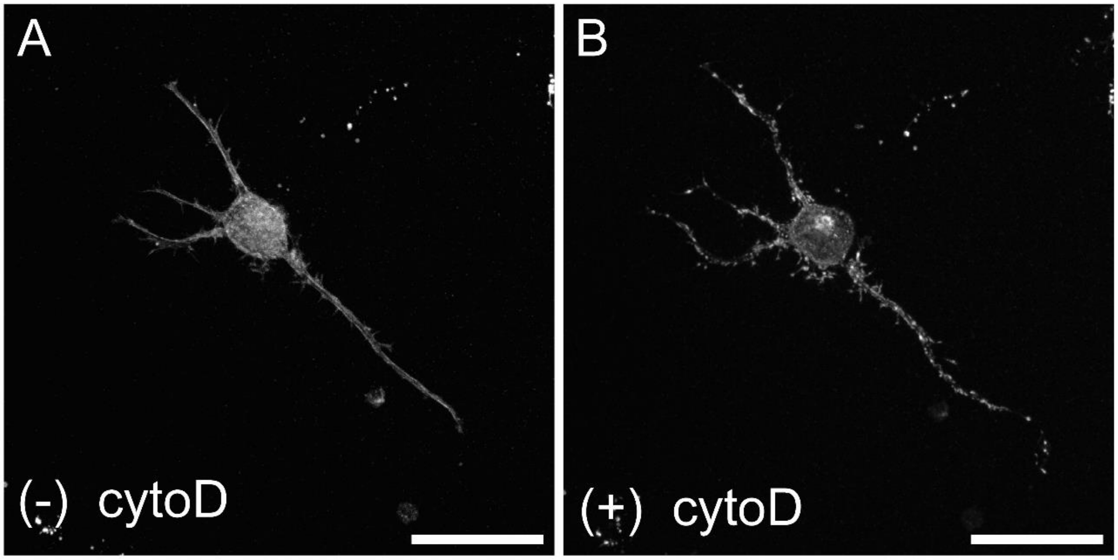


**Figure S17.** SPY650-FastAct live actin probe visualizing the organization of cellular actin (A) before and (B) after treatment with 4 uM cytochalasin D for 40 min. (scale bar = 50 μm)

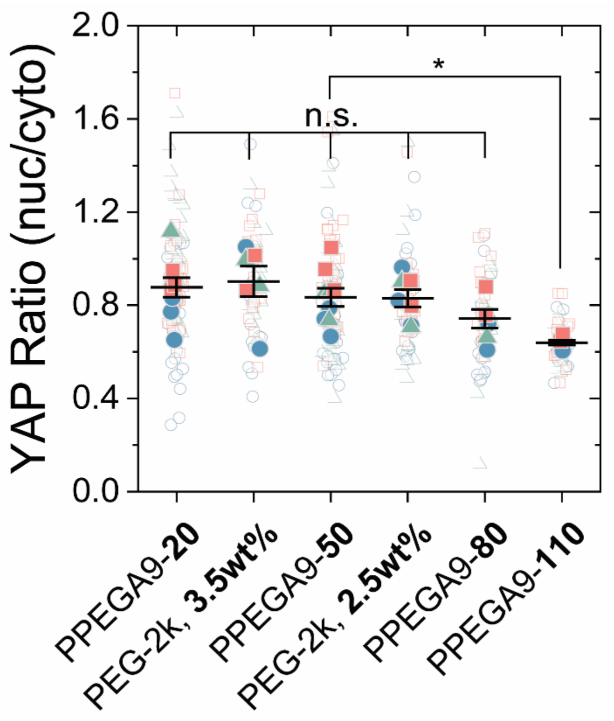


**Figure S18.** Measured nuclear/cytoplasmic YAP ratios measured for hMSCs cultured in each hydrogel formulation after 3 days of culture. (n = 3, independent biological replicates, at least 15 cells analyzed per replicate)

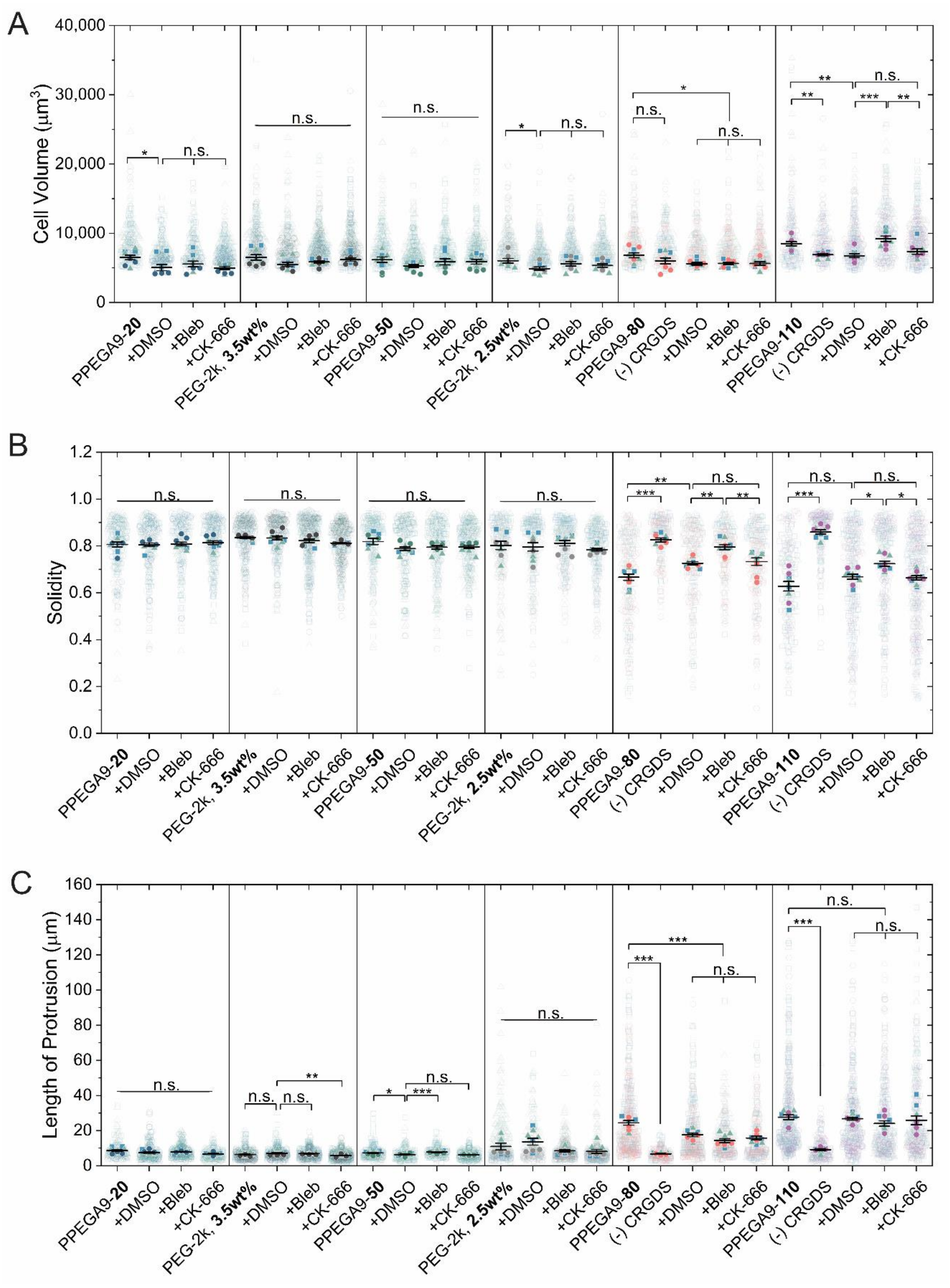


**Figure S19.** Summary super plots showing changes in (A) cell volume, (B) solidity, and (C) length of protrusions across all formulations and drug treatments. (n = 3, independent biological replicates, > 30 cells analyzed per replicate, mean ± SD analyzed using one-way ANOVA with Tukey's post hoc; **P*<0.05, ***P*> 0.01, ****P*>0.001)

**REFERENCES:**


[1] R. K. Das, V. Gocheva, R. Hammink, O. F. Zouani, A. E. Rowan, *Nat. Mater.* **2016**, *15*, 318.
[2] C. Storm, J. J. Pastore, F. C. MacKintosh, T. C. Lubensky, P. A. Janmey, *Nature* **2005**, *435*, 191.
